# Pleckstrin Homology-based proximity-labeling reveal lipid-associated interactomes

**DOI:** 10.64898/2026.11.12.765328

**Authors:** Lise C. Noack, Paul Vogel, Michaela Neubergerova, Roman Pleskot, Leonard Blaschek, Staffan Persson

**Affiliations:** Department of Plant & Environmental Sciences, Copenhagen Plant Science Center (CPSC), University of Copenhagen, Thorvaldsensvej 40, 1871 Frederiksberg, Denmark; Institute of Experimental Botany, Czech Academy of Sciences, Rozvojová 263, 165 00 Prague, Czech Republic; Department of Experimental Plant Biology, Faculty of Science, 12000 Charles University, Prague, Czech Republic; Umeå Plant Science Center, Department of Forest Genetics and Plant Physiology, Swedish University of Agricultural Sciences (SLU), 901 83 Umeå, Sweden; Joint International Research Laboratory of Metabolic & Developmental Sciences, State Key Laboratory of Hybrid Rice, School of Life Sciences and Biotechnology, Shanghai Jiao Tong University, Shanghai 200240, China

## Abstract

Phosphoinositides (PIPs) organize membrane-associated signaling, but the proteins associated with specific PIPs and their regulation remain poorly characterized. Here, we establish a lipid-binding domain-based proximity-labeling approach to map PIP-associated proteomes *in vivo*, and applied it to plant cells. We identify distinct protein networks associated with PI4P and PI(4,5)P₂, linking these lipids to membrane signaling, transport, and organization. Using molecular dynamics simulations and *in vivo* validation we further reveal how two previously uncharacterized PIP effectors, ALTERED RESPONSE TO GRAVITY-LIKE1 (ARL1) and HYPERSENSITIVE RESPONSE 1 (HIR1), engage with PIPs to regulate their membrane localization and nanodomain organization. Finally, we outline how PIP-associated proteomes change in response to prominent abiotic stress conditions, including salt and heat, in Arabidopsis. Our work establishes a platform for investigating lipid-associated proteomes *in vivo* and reveals how PIP-specific and stress-responsive protein networks contribute to membrane organization and cellular responses.

## Introduction

The plasma membrane (PM) harbors a diverse array of receptors, transporters, and signaling complexes, which are actively organized into specialized nanodomains^1,2^. Specific lipids, such as phosphoinositides (PIPs), sphingolipids, and sterols contribute to the formation of these nanodomains, which affect protein function and signal transduction efficiency.

PIPs are phospholipids with phosphorylated inositol ring headgroups that distribute differently across cellular membranes thus acting as molecular landmarks^3,4^. In particular, phosphatidylinositol-4,5-biphosphate (PI(4,5)P_2_) reside at the PM, while phosphatidylinositol-4-phosphate (PI4P) broadly localizes along the endocytic pathway, with an increasing gradient from the *trans*-Golgi Network/Early endosomes (TGN/EE) to the PM. PIPs interact with membrane-associated proteins via specific lipid-binding domains (LBDs) and motifs that are sensitive to membrane properties such as electrostaticity or membrane curvature^5–7^. Through these lipid-protein interactions, PIPs contribute to the spatial organization of signaling networks, as well as the regulation of protein conformation, stability and activity.

PIP conversion and degradation are rapid and dynamic as these processes are driven by a set of enzymes directly at the membrane. In Arabidopsis, phosphatidylinositol (PI) is converted into PI4P by the PI4 Kinases (PI4K) PI4Kβ1 and PI4Kβ2 at the TGN/EE, and PI4Kα1 at the PM^8,9^. In turn, PI4P can be converted into PI(4,5)P_2_ by PM-localized PI4P 5-kinase (PIP5K)^10^. The reverse conversion as well as degradation are mediated by PI-phosphatases and PI-phospholipases. Thus, PIP composition can change rapidly in response to environmental cues, such as salinity and temperature^11,12^, to promote membrane exchange. For example, salt-induced PI(4,5)P_2_ results in a general activation of endocytosis but also enlarged ER-PM connectivity^13,14^. Moreover, salt stress-induced PI4P activates PM H^+^-ATPases, and subsequently the SOS pathway^15^. Altogether, changes in PIPs enable rapid cellular responses to developmental cues and environmental stimuli.

Despite rapid advances in functional lipid probes, most methods aimed at identifying lipid-interacting proteins are not applicable to plant cells due to the presence of the cell wall^16^. In addition, prediction tools to identify protein domains that engage with PIPs are limited. Notably, LBD predictions, or electrostatic protein patches, are not enough to conclude *in vivo* interaction^5^. Therefore, identification of PIP effectors is particularly difficult, limiting our understanding of PIP regulatory mechanisms.

Recent advances in proximity-labeling to identify protein interactions have revolutionized protein complex identification. Of these labeling systems, biotin-based proximity labeling is most commonly used, e.g. TurboID and UltraID^17^. In addition, pupylation-based interaction tagging (PUP-IT) proximity labelling systems were recently adapted to plants and used to identify protein-protein interactions in signaling pathways and cell wall regulation^18–20^. Given the success in protein interaction identification, we hypothesized that proximity labeling may be exploited to also identify lipid-protein interactions.

Here, we developed a PUP-IT-based proximity-labeling approach to map PI4P- and PI(4,5)P₂-associated protein networks and uncovered how two previously uncharacterized effectors, ARL1 and HIR1, engage with PIPs to influence membrane localization and nanodomain organization. Finally, we show that these lipid–protein networks are dynamically remodeled under salt and heat stress, revealing the context-dependent nature of PIP-associated cellular processes.

## Results

### PI4P-LBD pupylation-based proximity labeling identifies a pool of membrane-associated proteins in Nicotiana benthamiana

To identify proteins localized to PIP-enriched membrane domains, we fused the PUP(E) ligase PafA to Lipid-Binding Domains (LBDs) **(Figure 1A)**. We first tested our approach using a specific LBD that targets PI4P; the Pleckstrin homology (PH) domain of Four-phosphate-adaptor protein1 (FAPP1), generating PafA-FAPP1, driven by a ubiquitin10 promoter (pUBQ10). In addition to binding PI4P, PH^FAPP^^1^ also binds the trans-Golgi network (TGN) resident protein ADP-ribosylation factor 1 (ARF1)^21^. To ensure target protein enrichment via solely PI4P, we introduced an E50A mutation in PH^FAPP^^1^ that prevents ARF1-binding^22^. In parallel, we generated two control constructs: a free GFP and a myristoylated and palmitoylated (MP)-mCherry fusion as general cytosolic and membrane controls, respectively **(Figure 1A).** Finally, we engineered the backbone to also hold a constitutively expressed FLAG-PupE fusion driven by a 35S promoter **(Figure S1A)**. Using these constructs, we confirmed efficient PUP-labeling of proteins via transient expression in *N. benthamiana* leaves where FLAG-labelled proteins were precipitated by anti-FLAG immunoblotting prior to identification and quantification by label-free mass spectrometry **(Figure S1B, Supplementary Dataset 1)**.

**Figure 1.**
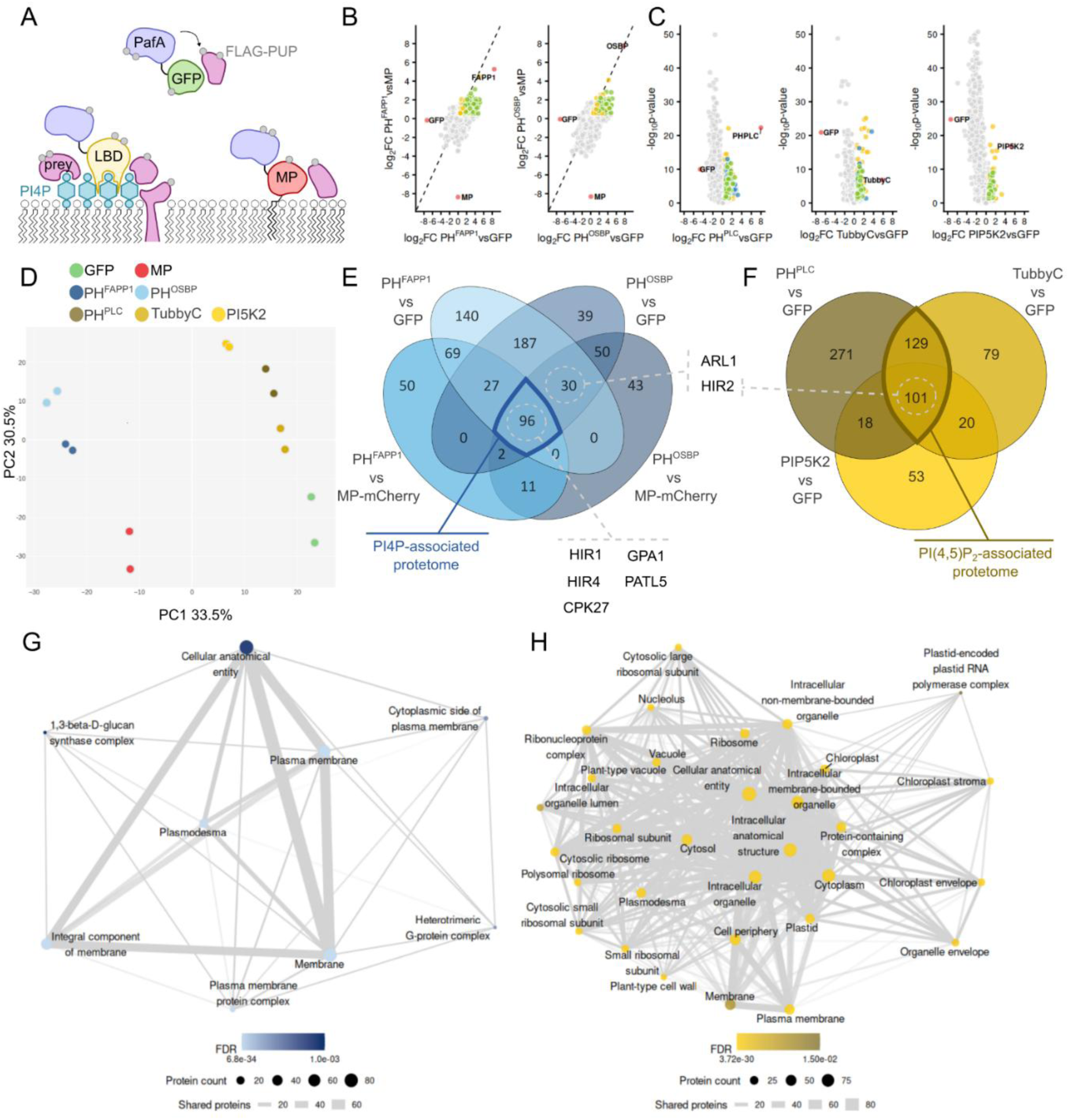
Identification of the PIP-associated proteomes by Pupylation-Based Proximity Labeling in A. thaliana. **A.** Illustration of the PUP-IT system used in this study. PafA fused to the Lipid-Binding Domain (LBD) labels proteins found in regions rich in the corresponding phosphoinositide, while a GFP fusion and a MP-mCherry (MP) act as cytosolic and membrane controls for nonspecific enrichment. **B.** Volcano plots showing proteins significantly enriched in PH^FAPP1^ and PH^OSBP^ versus controls in yellow. Proteins enriched in both PH^FAPP1^ and PH^OSBP^ are shown in green. Baits are shown in red. **C.** Volcano plots showing proteins significantly enriched in PH^PLC^, TubbyC and PIP5K2 versus controls in yellow. Proteins enriched in both PH^PLC^ and TubbyC are shown in blue, while proteins enriched in PH^PLC^, TubbyC and PIP5K2 are shown in green. Baits are shown in red. **D.** Principal component analysis on normalized protein intensity of PUP-IT constructs expressed in Arabidopsis thaliana seedlings. **E,F.** Venn Diagram of the proteins found with a log_2_FC>0.5 and an adjusted p-value>0.05 in the different comparisons. **G,H.** Gene Ontology analysis of cellular compartment for proteins enriched in the PI4P- and PI(4,5)P_2_-associated proteomes.

Principal component analysis (PCA) on mean protein intensity showed that PafA-PH^FAPP^^1^ samples clustered separately from the membrane control, indicating that PafA-PH^FAPP^^1^ and MP-mCherry labeled different pools of proteins despite residing in similar cell compartments **(Figure S1C)**. From the MS-analyses, we confirmed the strong labeling of the baits **(Figure S1E)** and identified 320 proteins enriched in PafA-PH^FAPP1^ vs controls **(Figure S1D and E, Supplementary Dataset 2)** including PM or endomembrane system, in line with the presence of PI4P in those compartments **(Figure S1E)**. In addition, several proteins known to interact with, or be regulated by, PIPs were identified confirming that our method is suitable to identify proteins associated with PI4P-enriched membranes in *N. benthamiana* leaves **(Figure S1E, Table Supp 1)**.

### Identification of the PIP-interactomes by Pupylation-Based Proximity Labeling in Arabidopsis

As our results from heterologous expression of PH^FAPP1^ baits in *N. benthamiana* were promising, we decided to map proteins associated with PIP-enriched membranes in Arabidopsis seedlings. To do so, we transformed the PUP-IT constructs stably into Arabidopsis wild-type plants. In addition, to limit the risk of identifying interactions specifically mediated by PH^FAPP1^ we included the PH domain of the Oxysterol-Binding protein (PH^OSBP^) as a second PI4P-specific bait **(Figure S2A)**. Additionally, we extended our approach to include PI(4,5)P_2_, using two LBDs – the PH domain of phospholipase C (PH^PLC^) and the C-terminal domain of the TUBBY protein (TubbyC) – as well as the Phosphatidyl-inositol Kinase PIP5K2, which phosphorylates PI4P to generate PI(4,5)P_2_ at the PM. We confirmed efficient PUP-labeling of proteins by western blotting after FLAG enrichment in the corresponding transgenic lines **(Figure S2B)**. Notably, we did not see any deviation in growth or development, aside from some slight delay in root growth of some of the Arabidopsis PUP-IT lines, compared to wild type (**Figure S2C and D)**. We therefore used the FLAG-enriched samples for protein identification and quantification by label-free mass spectrometry, from which we confirmed strong labeling of the baits **(Figure 1B and C)**. Finally, PCA on mean protein intensity revealed that proteins identified by the PI4P baits clustered together, as did the proteins identified by the three PI(4,5)P_2_ baits. By contrast, the PI4P enriched proteins clearly separated from the PI(4,5)P_2_ enriched ones, indicating labelling of specific protein pools for each PIP species **(Figure 1D, Supplementary Dataset 3)**.

### PI4P-interactome shows an enrichment in receptors and transporter activity

We identified a total of 96 enriched proteins shared between PH^FAPP1^ and PH^OSBP^ compared to controls. These 96 proteins (referred to as the PI4P-associated proteome) corresponded to 50% of the total PH^FAPP1^ and 55% of the total PH^OSBP^-enriched proteins **(Figure 1B, 1E and S2C, Supplementary Dataset 4)**. We first performed a Gene Ontology (GO) term enrichment analysis on cellular compartment, which confirmed that the PI4P-associated proteome predominantly contained proteins from the PM and the endomembrane system **(Figure 1G, Supplementary Dataset 5)**. This result is concordant with previous work demonstrating that the major pool of PI4P is localized at the PM, while a minor pool is present at the TGN/EE^22^. Several known interactors of PIPs were among the PI4P-associated proteome confirming that our approach is suitable to find PIP-associated proteins in Arabidopsis (**Figure S3A, Table Supp 2**).

While we did not find any significant enrichment in lipidated proteins and proteins containing PIP-interacting motifs (e.g. C2, SEC14, PH or ENTH domains) in the PI4P-associated proteome compared to the membrane proteome, i.e. proteins enriched in MP-mCherry bait relative to the GFP bait **(Figure S4A and B**), we noted a significant enrichment in transmembrane proteins, representing over 50% of proteins **(Figure S3C).** Specifically, protein kinases and ABC transporters were the two main protein families in the PI4P-associated proteome **(Figure S4D)**. The predominance of signaling function in the PI4P-associated proteome was further supported by the GO term enrichment in “Phosphorylation” together with “G-coupled receptor signaling pathway” **(Figure S4E)**. In addition, “Callose synthesis” and “Vesicular transport” were enriched in the PI4P-associated proteome.

### The PI(4,5)P_2_-associated proteome was enriched in transport function

PI(4,5)P_2_ is present in low concentration in membranes, reflected in a strong cytosolic signal of fluorescently-tagged PI(4,5)P_2_ LBDs^23^,and of PI5K2 fluorescent reporters^24^. As the MP-mCherry control is strongly associated with the PM with little cytosolic distribution, we therefore argued that the free GFP is a more appropriate control for the PI(4,5)P_2_-related proximity labelling experiments. Using this set-up, we found 230 proteins (referred to as the PI(4,5)P_2_-associated proteome) enriched by both LBDs (i.e. PLC and TubbyC), representing 44% and 70% of total PH^PLC^-enriched and TubbyC-enriched proteins, respectively **(Figure 1C, 1F and S3B, Table 5)**. Further including the PI5K-enriched proteins, the PI(4,5)P_2_-associated proteome could be refined to 101 overlapping proteins, representing 19%, 30%, and 52% of total PH^PLC^, TubbyC and PI5K2-enriched proteins, respectively.

The PI(4,5)P_2_-associated proteome comprised PM and endomembrane resident proteins, including some known PIP effectors **(Table Supp 3)**. Several cytosolic, plastid-localized and ribosomal proteins were also found. Although we cannot exclude that this enrichment is biologically relevant, it could represent non-specific labelling by the baits as PI(4,5)P_2_ is present only at very low levels in membranes under control conditions **(Figure 1H)**. Nevertheless, GO term analysis revealed an enrichment of “Export across the membrane” with three PM H^+^ ATPases (AHA1, AHA2 and AHA11) and 5 aquaporins (PIP1-1, PIP2-1, PIP2-2, PIP2-3 and PIP2-7) in the PI(4,5)P_2_-interactome **(Figure S4 F, Supplementary Dataset 4 and 5)**.

### PI4P-interacting proteins change their distribution patterns and dynamics upon PIP depletion

To validate the data obtained from the PI4P PUP-IT interactome, we selected several candidate proteins enriched in the PI4P-associated proteome for further studies **(Figure 1E)**. These included the alpha subunit of heterotrimeric G protein (GPA1), the sec14 containing protein Patellin 5 (PATL5), the Calcium-dependent Protein Kinase 27 (CPK27), the Altered response to Gravity-Like1 (ARL1), BAK1-INTERACTING RECEPTOR-LIKE KINASE 2 (BIR2) and HYPERSENSITIVE RESPONSE 1 (HIR1). BIR2 is a transmembrane helix-containing protein, while GPA1, CPK27 and HIR1 are lipidated^25–28^. Finally, PATL5 and ARL1 are membrane-associated proteins without lipid anchors^29,30^. We first confirmed membrane localization of these proteins by confocal microscopy in root epidermal cells of Arabidopsis. As previously reported, BIR2 was exclusively present at the PM^25,31^. ARL1 and PATL5 were localized at the PM, as well as in the cytosol similarly to the reported localization of close homologs ARL2 and PATL3, respectively **(Figure 2A)**^30,32^. We further confirmed dual localization of GPA1 at the PM and at the TGN/EE^33^, and that HIR1 was localized at the PM but also showed faint labelling of the tonoplast **(Figure 2C)**^28^. Finally, we confirmed the PM localization of CPK27, previously reported in protoplasts, and additionally noticed the presence of a faint signal at the tonoplast **(Figure 2A)**^26^.

**Figure 2.**
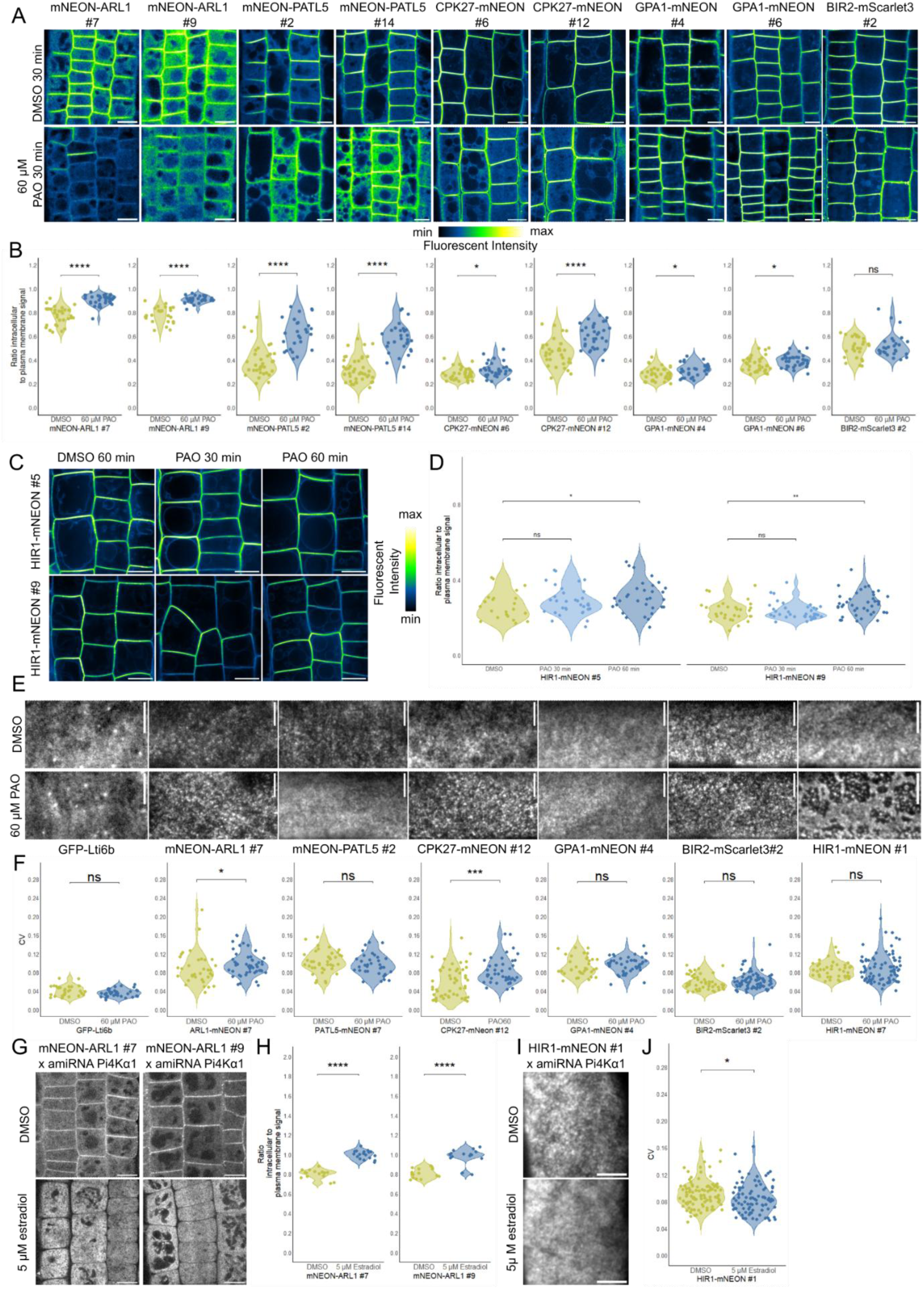
PI4P-associated protein candidates change their distribution patterns and dynamics upon PIP depletion A,B. Confocal microscopy images (A) and calculated ratio of intracellular to plasma membrane fluorescent signal (B) of Arabidopsis root epidermal cells expressing pUBQ10:mNeon-ARL1, pUBQ10:mNeon-PATL5, pUBQ10:CPK27-mNeon, pUBQ10:GPA1-mNeon, and pUBQ10:BIR2-mScarlet3 treated with 60 µM PAO for 30 min or the equivalent volume of DMSO. Scale bar = 10 µm. Student’s T test was used to assess statistical significance when normal distribution was confirmed using the Shapiro test. Wilcoxon rank-sum tests were used otherwise. (n ≥ 24). **C,D.** Confocal microscopy images (C) and calculated ratio of intracellular to plasma membrane fluorescent signal (D) of *Arabidopsis* root epidermal cells expressing p35S:HIR1-mNeon, treated with 60 µM PAO for 30 min or 60 min or the equivalent volume of DMSO for 60 min. Scale bar = 10 µm. Wilcoxon rank-sum tests were used for statistical analysis. (n ≥ 23). **E,F.** TIRF microscopy images (E) and coefficient of variation (CV) of fluorescent signal along a 10µm line (G) of *Arabidopsis* root epidermal cells expressing p35S:GFP-Lti6b, pUBQ10:mNeon-ARL1, pUBQ10:mNeon-PATL5, pUBQ10:CPK27-mNeon, pUBQ10:GPA1-mNeon, pUBQ10:BIR2-mScarlet3, and p35S:HIR1-mNeon, treated with 60 µM PAO or the equivalent volume of DMSO for 30 min. Scale bar = 5µm. Student’s T test was used to assess statistical significance when normal distribution was confirmed using the Shapiro test. Wilcoxon rank-sum tests were used otherwise. (n ≥ 29). **G,H.** Confocal microscopy images (G) and calculated ratio of intracellular to plasma membrane fluorescent signal (H) of *Arabidopsis* root epidermal cells expressing pUBQ10:mNeon-ARL1 carrying the amiRNA_PI4Kα1 treated with 5 µM β-estradiol or the equivalent volume of DMSO. Scale bar = 10 µm. Student’s T test was used to assess statistical significance when normal distribution was confirmed using the Shapiro test. Wilcoxon rank-sum tests were used otherwise. (n ≥ 15). **I,J.** TIRF microscopy images (I) and coefficient of variation (CV) of fluorescent signal along a 10µm line (J) of *Arabidopsis* root epidermal cells expressing p35S:HIR1-mNeon carrying the amiRNA_PI4Kα1 treated with 5 µM β-estradiol or the equivalent volume of DMSO. Scale bar = 5 µm. Wilcoxon rank-sum test was used. (n ≥ 92).

We then tested how the protein localization was affected upon reduction in PIPs using the PIP Kinase inhibitor PAO (60µM) for 30min, which affects both PI4P and PI(4,5)P_2_ levels at the PM of root epidermal cells **(Figure 2A)**^22^. PAO increased the ratio of intracellular to PM fluorescent signal for GPA1, CPK27, PATL5 and ARL1 **(Figure 2B)**. No changes were observed for HIR1 after 30min, but longer treatment (60min) showed an increased ratio of intracellular to PM fluorescence **(Figure 2C and 2D).** In the case of CPK27, PATL5 and ARL1, the changes in ratio were mostly driven by a reduced PM signal, indicating that PIPs are necessary for their PM localization **(Figure S5A,B)**. By contrast, PAO treatment did not affect the GPA1 signal at PM, but instead drastically reduced its endomembrane localization **(Figure S5A-D)**, indicating that the PI4P pool at the TGN regulates GPA1 subcellular localization. Finally, BIR2 fluorescent signal at the PM was not affected, but PIP inhibition led to enriched BIR2 signal in endomembrane compartment, suggesting that BIR2 vesicular trafficking was impacted **(Figure 2A and S5A,B)**.

To gain a better understanding of the effect of PIP depletion on protein dynamic, we performed Total Internal Reflection Fluorescence (TIRF) microscopy upon PAO treatment for the selected candidate proteins, as well as Lti6b as a membrane protein control. Lti6b, PATL5, GPA1 and BIR2 signal distribution at the PM did not show any obvious response to PAO according to the calculated coefficient of variation (CV) of fluorescent signal that measures the heterogeneity of signal distribution at the PM **(Figure 2E and F)**. However, we observed an increase in ARL1 and CPK27 nanodomain formation upon PAO treatment. Surprisingly, HIR1 signal showed reticulate-like patterns at the PM, distinctly different from the DMSO control. Similar patterns were not observed for any of the other membrane proteins suggesting a HIR-specific response to PAO. Based on these results, we hypothesize that HIR1, ARL1 and CPK27 might engage with PIPs at the PM and that this interaction affects the nanodomain organization or stability.

As PATL5 and ARL1 are peripheral membrane proteins, we tested how PI4P depletion affected their localization using *Nicotiana benthamiana* SAC1 assays^34^. In this assay, PATL5 and ARL1 were co-expressed with the PI4-phosphatase SAC1 targeted to the PM via a membrane anchor. PI4P depletion by SAC1 resulted in a delocalization of both PATL5 and ARL1 in the cytosol, showing that ARL1 and PATL5 rely specifically on PI4P **(Figure S5E).** Taken together, PIP alterations affected the distribution of several proteins enriched in our PI4P-enriched interactome.

### PI4Kα1-dependent PI4P controls ARL1 localization and HIR1 nanodomain stability

We selected ARL1 and HIR1 for further investigation of their membrane-binding mechanisms, as both responded strongly to PIP alterations and were affected in their membrane binding and nanodomain formation, respectively. We first took advantage of the β-estradiol inducible artificial microRNA targeting PI4Kα1 (amiRNA PI4Kα1) to test the effect of a deregulation of the PM pool of PI4P on ARL1 and HIR1^9^. amiRNA PI4Kα1 induction resulted in a delocalization of ARL1 to the cytosol, similar to what we observed with PAO treatment, but did not affect HIR1 localization **(Figure 2 G and H, Figure S5 F and G)**. However, using Brefeldin A (BFA) to aggregate endosomes, TGN and ER into BFA bodies, we observed that PI4Kα1 knockdown resulted in a stronger HIR1 fluorescent signal in BFA bodies indicating a higher proportion of HIR1 proteins present in intracellular compartment **(Figure S5H and I)**. Furthermore, using TIRF microscopy, we observed that PI4Kα1 knockdown affected the HIR1 nanodomain organization as indicated by a lower CV **(Figure 2I and J)**. As HIR1 is not only present at the PM but also in intracellular compartment, we also crossed HIR1 reporter lines with the double mutant *pi4kβ1pi4kβ2* but we did not observe a delocalization of HIR1 **(Figure S5 J and K)**. These results indicated that the pool of PI4P produced by PI4Kα1 at the plasma membrane controls ARL1 localization and HIR1 nanodomain stability at the PM.

### Molecular dynamics simulations indicate ARL1 dimerization with preferential binding to PI4P

ARL1 is predicted to form homodimers **(Figure S6A)**, with the globular N-termini forming a positively charged surface, which could interact with negatively charged membranes **(Figure S6B)**. To investigate this, we performed three independent replicas of a 10µs coarse grained molecular simulation of the ARL1 dimer (1-354), excluding the disordered C-terminal region. We positioned the dimer close to a complex membrane made of phosphatidyl choline (PC) and phosphatidylethanolamine (PE), as well as negatively charged lipids: PI4P (PIP1), PI(4,5)P_2_ (PIP2), phosphatidylserine (PS) and phosphatidic acid (PA) **(Figure 3A)**. In all simulations, the ARL1 dimer stably bound the membrane through the N-terminal positively charged surface. In addition, ARL1 showed a preferential binding for PI4P and PI(4,5)P_2_ over PA and PS **(Figure 3B,C and S6D,E)**.

**Figure 3.**
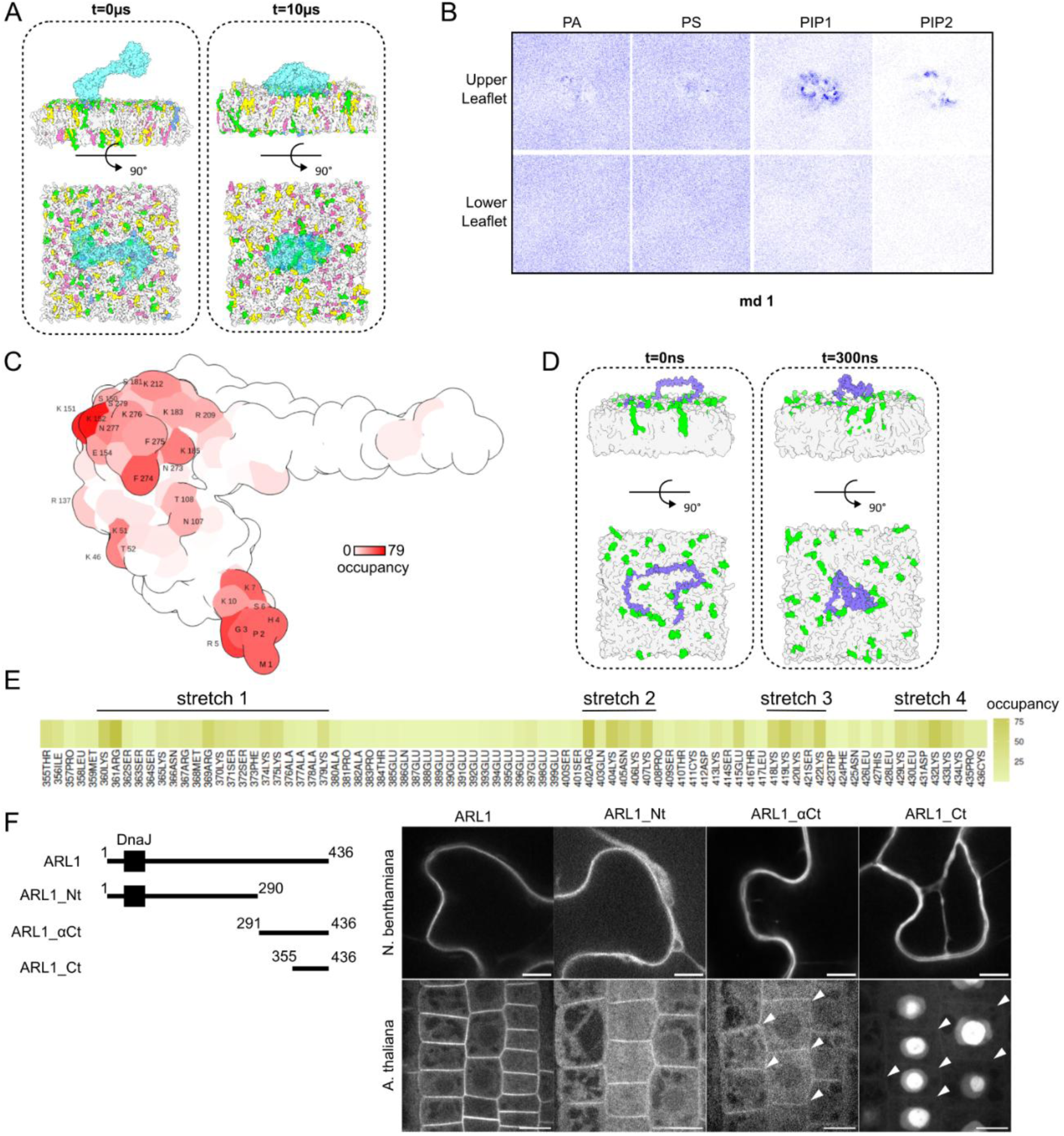
The ARL1 dimer binds the plasma membrane through its N-terminal domain. **A.** CG-MD simulations of ARL1(1-354) dimer with a complex negatively charged membrane containing 1% PIP2 (in blue), 5% PIP1 (in green), 10% PA (in pink), 10% PS (in yellow), 37% PC (in grey) and 33% PE (in grey). ARL1 is colored in cyan. The initial conditions are shown on the left side (0 µs) and the membrane-bound protein complex on the right side (10 µs). **B.** Two-dimensional density (over 10 µs) of PIP1, PIP2, PA and PS in upper leaflet (adjacent to the protein) and lower leaflet. **C.** Occupancy map of PIP1 on ARL1(1-354) structural representation. The occupancy is defined as the percentage of frame with contact (distance under 0.7 Å). **D.** Atomistic-MD simulations of the ARL1(355-end) C-terminal disorder region (in purple) with a POPC membrane (in grey) containing 10% PI4P in the leaflet adjavent to the protein (in green). The initial conditions are shown on the left side (0 ns) and the membrane-bound protein on the right side (300 ns). **E.** Occupancy of PI4P at each amino acids of the ARL1(355-end) C-terminal disordered region. **F.** Confocal microscopy images of Nicotiana leaf epidermal cells (upper panel) and *Arabidopsis* root epidermal cells expressing mNEON-ARL1 full length, ARL1(1-290)-mNEON (ARL1_Nt), mNeon-ARL1(291-436) (ARL1_αCt) or mNEON-ARL1(355-436) (ARL1_Ct). Faint plasma membrane signal is highlighted with white arrows. Scale bar = 20 µm in upper panel and 10 µm in lower panel.

The disordered C-terminus region of ARL1 carries four stretches of positively charged amino acids, interrupted by a stretch of glutamic acids and could also contribute to the binding of ARL1 to membranes **(Figure 3E)**. Atomistic simulation of the disordered C-terminal domain (355-end) with a membrane composed of PC containing 10% PI4P revealed that the ARL1 C-terminal domain stayed bound to the membrane through the different stretches of positively charged amino acids while it curls up around the stretch of glutamic acids **(Figure 3D,E)**.

Our model therefore indicated that the ARL1 dimer could interact with the membrane via both the N-terminal surface and C-terminal stretches. We tested this hypothesis *in planta* by expressing truncated versions of ARL1 transiently in *N. benthamiana* leaves and stably in *Arabidopsis* **(Figure 3F)**. The ARL1 (1-290) N-terminus was sufficient to localize at the PM, and the ARL1(355-end) C-terminus alone shows a faint signal at the PM. The addition of the alpha helix (291-436) was sufficient to partially restore localization to the PM. This might be due to the alpha helix allowing dimerization with native full length ARL1 proteins. We thus conclude that the N-terminal surface is the main membrane-binding domain of ARL1, with the possibility that the C-terminus stabilizes or orients the protein at the membrane.

### Molecular dynamics simulations indicate how HIR proteins engage with PIPs

HIR1 belongs to the HYPERSENSITIVE RESPONSE (HIR) protein family, a plant-specific family of stomatin/prohibitin/flotillin/HflK/C (SPFH) proteins composed of four members, of which three mainly localized at the PM: HIR1, HIR2 and HIR4^28,35,36^. Interestingly, we found HIR1 and HIR4 in the PI4P-associated proteome, while HIR2 was enriched in PH^OSBP^ as well as in the PI(4,5)P_2_-associated proteome indicating that the association of HIR protein with PIPs is not restricted to HIR1 **(Figure 1E)**. To first understand how the HIR proteins might engage with PIPs, we generated a coarse-grained representation of a HIR1 complex composed of 14 subunits **(Figure 4A)**. The number of subunits was based on the iPTM values obtained from AlphaFold 3 prediction and is of similar order of magnitude to what has been previously observed for HIR2 **(Figure S7A and B)**^31,36^. The N-termini of HIR1 14-mer form a hydrophobic ring in contact with the membrane and are overlooked by positively charged amino acids **(Figure S7C and D)**. HIR1 carries myristoylation and palmitoylation sites at position G2 and C6, respectively^37,38^. We thus added these modifications before positioning the HIR1 complex on a membrane containing equal amount of PE and PC and 1% PI(4,5)P_2_, 5% PI4P, 10% PA, and 10% PS in the cytosolic leaflet in direct contact with the protein complex **(Figure 4A)**.

**Figure 4.**
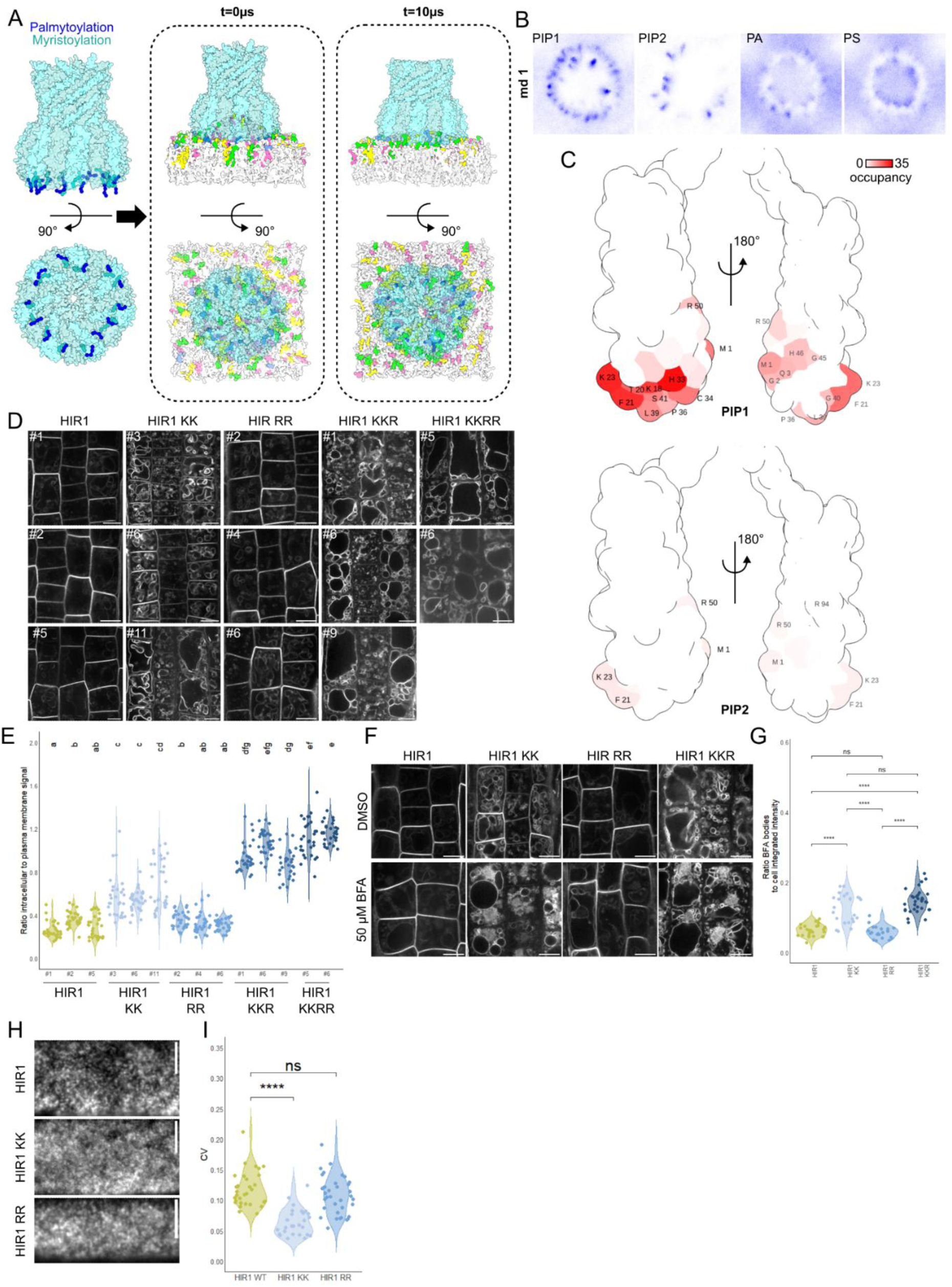
HIR1 oligomer binds PIPs via positively charged amino acids which are necessary for HIR1 stability at the PM. **A.** CG-MD simulations of the HIR1 complex composed of 14 subunits carrying myristoylation in G2 and palmitoylation in C7 with a complex negatively charged membrane containing 1% PIP2 (in blue), 5% PIP1 (in green), 10% PA (in pink), 10% PS (in yellow), 37% PC (in grey) and 33% PE (in grey) in the cytosolic leaflet and 50% PC, 50% PE in the extracellular leaflet. HIR1 complex is colored in cyan. The initial and final conditions are shown on the left side (0 µs) and right side (10 µs), respectively. **B.** Two-dimensional density (over 10 µs) of the different lipid molecules in the cytosolic leaflet adjacent to the HIR1 complex. **C.** Occupancy map of PIP1 and PIP2 on the HIR1 SPFH domain. The occupancy is defined as the % of frame with contact (distance under 0.7 Å). **D.** Confocal microscopy images of Arabidopsis root epidermal cells expressing p35S:HIR1-mNEON (HIR1), p35S:HIR1(K18Q;K23Q)-mNeon (HIR1 KK), p35S:HIR1(R50Q;R94Q)-mNeon (HIR1 RR), p35S:HIR1(K18Q;K23Q;R50Q)-mNeon (HIR1 KKR), p35S:HIR1(K18Q;K23Q;R50Q;R94Q)-mNeon (HIR1 KKRR). Scale bar = 10 µm **E.** Ratio of intracellular to plasma membrane fluorescent signal in epidermal root cells of HIR1 reporter lines. Statistical analysis was done using Krustal-Wallis’s test, followed by Dunn’s test with Benjamini-Hochberg correction. (n ≥ 30) **F.** Confocal microscopy images of *Arabidopsis* root epidermal cells expressing p35S:HIR1-mNEON (HIR1), p35S:HIR1(K18Q;K23Q)-mNeon (HIR1 KK), p35S:HIR1(R50Q;R94Q)-mNeon (HIR1 RR), and p35S:HIR1(K18Q;K23Q;R50Q)-mNeon (HIR1 KKR) treated with 50 µM BFA for 60 min or the equivalent volume of DMSO. Scale bar = 10 µm **G.** Ratio of integrated intensity of BFA bodies to the whole cell in HIR1 reporter lines treated with 50 µM BFA for 60 min. Statistical analysis was done using a pairwise comparisons Wilcoxon rank-sum test. (n ≥ 22). **H.** TIRF microscopy images of Arabidopsis root epidermal cells expressing p35S:HIR1-mNeon and p35S:HIR1(K18Q;K23Q)-mNeon (HIR1 KK). Scale bar = 5µm **I.** Quantification of the coefficient of variation (CV) of fluorescent signal along a 10 µm line. Statistical analysis was done using a pairwise comparisons Wilcoxon rank-sum test. (n ≥ 27)

We found that PI4P and PI(4,5)P_2_ clustered around the HIR1 complex more than PA and PS, in all performed simulation (n=5) **(Figure 4B, C and S7E)**, likely mediated via two different interaction interfaces. The first one, on the outside of the HIR1 complex, is composed of the positively charged amino acids K18, K23 and H33 **(Figure 4C)**. The second one is situated on the inner side of the HIR complex and is formed by the positively charged amino acids R50 and H46. We also noted that PI(4,5)P_2_ was able to interact with R94.

To refine our model, we performed atomistic molecular dynamics simulations of the HIR1 SPFH domain (1-154) with a membrane of PC containing 10% PI4P in each leaflet. The simulation confirmed the binding of PI4P to HIR1 predominantly via interaction with K23 and R50 **(Figure S7 G and H)**. Consistent with these results, the mutations K18Q, K23Q and R50Q in HIR1 were sufficient to suppress PI4P binding in the atomistic simulations.

### Distinct amino acids in HIR1 underpin its PM interactions and nanodomain organization

To investigate whether the predicted amino acids in the molecular dynamics simulations contribute to the associations of HIR proteins and PIPs, we made HIR1 KKR variants in which we mutated K18, K23, and R50 to glutamate (HIR1 KKR) or added a mutation in R94 (HIR1 KKRR). We also generated a K18QK23Q variant where only the outer interface was mutated (HIR1 KK) as well as a R50Q mutant (HIR1 R) and a R50QR94Q mutant (HIR1 RR) in which only the inner interface was mutated. We fused these variants to mNeonGreen (mNeon) and transiently expressed them in *N. benthamiana* leaves. Here, HIR1 mainly localized to the PM **(Figure S8A)**, while mutations of the outer interface (KK) and inner and outer interfaces (KKR or KKRR) altered the localization to mainly endomembrane compartments **(Figure S8A)**. Using wave marker lines for endomembrane compartments, we observed that the increased labelling of HIR1 KKR compartments coincided with the TGN marker primarily and Golgi secondarily **(Figure S8C and D)**^39^. By contrast, mutation of R50 only at the inner interface did not affect HIR1 localization.

We generated Arabidopsis lines with the same HIR1 variants, apart from HIR1 R50Q as no lines with fluorescence could be retrieved. Mutations of the outer interface (KK) resulted in accumulation of the protein at the tonoplast, while mutation in the inner interface (HIR1 RR) did not affect HIR1 localization **(Figure 4D and E)**. Similar to the observations in *N. benthamiana*, combined mutations of the inner and outer interfaces (KKR and KKRR) had a synergistic effect, resulting in strong tonoplast labelling and little to no signal at the PM. To evaluate if some fluorescence might also accumulate in other endomembrane compartments, we used BFA which revealed that a higher proportion of HIR1 proteins were present in trafficking compartments in the KK and KKR lines than in the HIR1 WT or RR lines **(Figure 5 F and G)**. Taken together, these results indicate that the interaction between HIR1 and PIPs is largely maintained via the outer interface of HIR1, and that distinct amino acids in this region of HIR1 are necessary for the PM localization.

**Figure 5.**
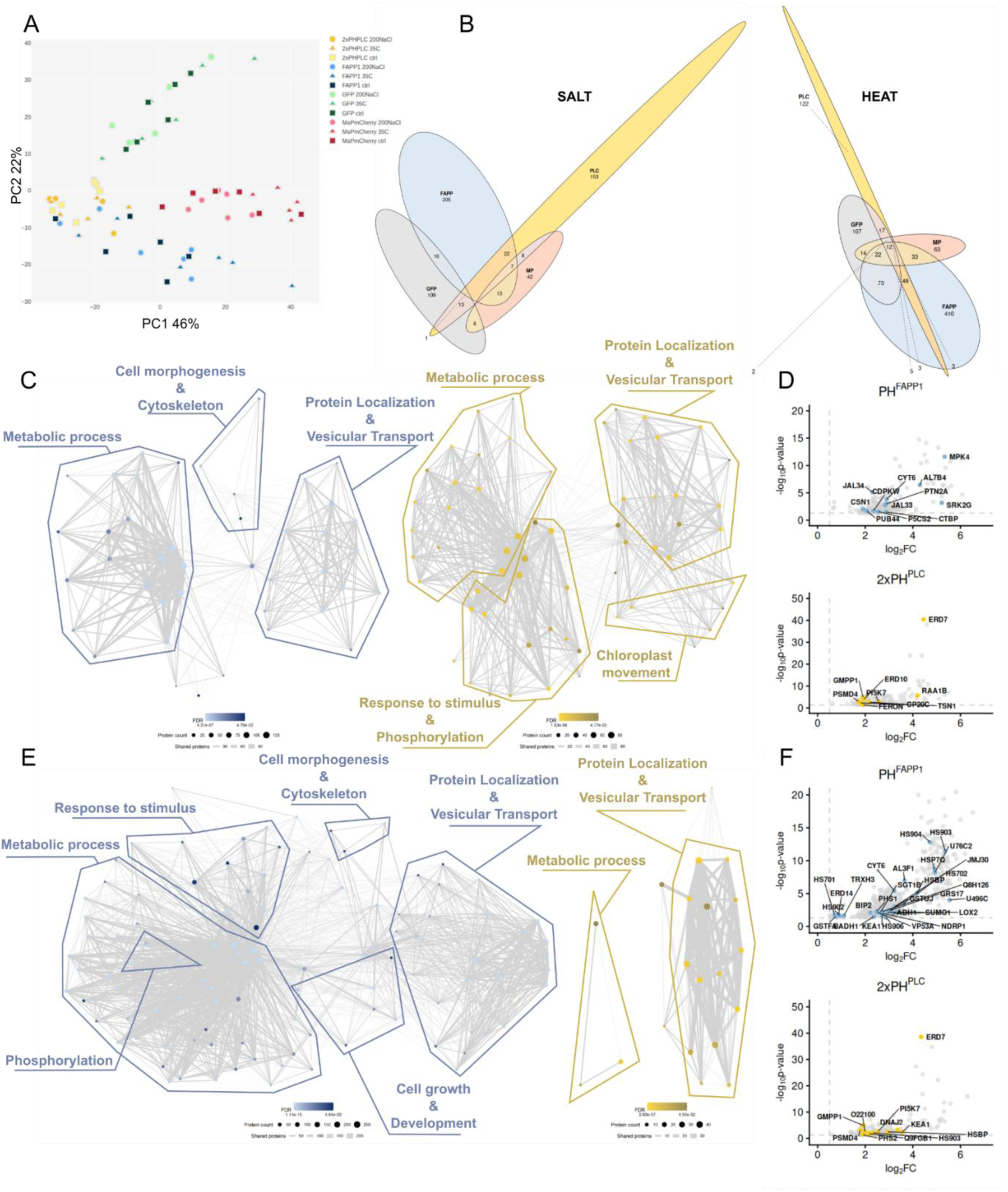
PI4P and PI(4,5)P_2_-associated proteomes respond differently to abiotic stresses. **A.** Principal component analysis on normalized protein intensity of estradiol-inducible PUP-IT constructs expressed in *Arabidopsis thaliana* seedlings in control conditions, upon salt stress (200mM NaCl) or upon heat stress (35°C) for 2h. **B.** Venn Diagram of proteins significantly enriched in salt stress (right panel) and heat stress (left panel) compared to control conditions for GFP, MP-mCherry, PH^FAPP1^, and 2xPH^PLC^ baits. **C, E.** Gene Ontology analysis of cellular processes for proteins enriched in salt stress (C) and heat stress (E) compared to control conditions for PH^FAPP1^ in blue and 2xPH^PLC^ in yellow. Clusters of GO terms have been annotated based on the main cellular process. D, F. Volcano Plot for proteins enriched in salt stress (D) and heat stress (F) compared to control conditions for PH^FAPP1^ in blue and 2xPH^PLC^ in yellow. Proteins associated with salt (D) and heat stress (F) are annotated.

We next assessed how PIP perturbation impacted HIR1 organization at the PM using TIRF microscopy to first image transiently expressed HIR1 and its variants in *N. benthamiana* leaves. Here, the mutated variants showed a minor, but significant decrease in CV of the signal, meaning a more homogeneous signal at the PM for HIR1 R, KKR and KKRR **(Figure S8A and B)**. Similarly, in *A. thaliana*, the variants with mutations in the outer interface (KK) led to a lower CV **(Figure 4H and I)**. However, mutations in the inner interface (RR) did not significantly affect nanodomain organization, suggesting once again a prevalent role of the PIPs interaction on the outer interface of the HIR1 complex.

### PI4P and PI(4,5)P_2_-associated proteomes respond differently to abiotic stresses

As PIPs are important for responses to different abiotic stresses, we assessed how the PIP-associated proteomes change in response to salt and heat stress. While the PI(4,5)P_2_ content at the PM is low under normal conditions, it rapidly increases in response to salt and heat stress^11,12^. Thus, under those conditions, labeling of PI(4,5)P_2_ effectors might be more prominent. To adapt our system to the study of stress response, we modified the expression of the substrate FLAG-PupE to be under the control of an estradiol-inducible promoter and introduced it into Arabidopsis together with PafA-2xPH^PLC^ and PafA-PH^FAPP1^ to follow PI(4,5)P_2_ and PI4P-associated proteomes, respectively, as well as MP-mCherry-PafA as control **(Figure S9A)**. We tested the PUP-labelling at different time points by western blot, where most lines showed substantial labelling signal with 6 to 8 h estradiol-induction **(Figure S9B)**. To ensure sufficient time for PUP-IT induction, we induced the plants for 6 h, followed by a 2 h treatment with 200 mM NaCl for salt stress or 2 h heat stress at 35°C. The experiment was reproduced twice, and the results were merged to identify proteins enriched consistently over both experiments. After enrichment and quantification by label-free mass spectrometry, we confirmed the presence of PupE-FLAG and PafA proteins in every sample, indicative of efficient labelling **(Figure S9C, Supplementary Dataset 6)**.

The results of PCA on mean protein intensity showed that PI4P and PI(4,5)P_2_ baits clustered separately from the control baits under control and stress conditions. In addition, PI4P and PI(4,5)P_2_ baits separated with a slight overlap indicating labelling of both a common pool of proteins as well as a specific protein pool for each PIP **(Figure 5A)**. More specifically, we found 205 and 153 proteins enriched for PH^FAPP1^ and 2xPH^PLC^, respectively, during salt stress and 410 and 122 enriched in PH^FAPP1^ and 2xPH^PLC^, respectively, upon heat stress **(Figure 5B, Supplementary Dataset 7-8)**. In addition, 22 and 48 proteins were enriched in both PH^FAPP1^ and 2xPH^PLC^, respectively, under salt and heat stress, highlighting the overlap between the two baits. To evaluate if the enrichment observed was solely due to transcriptional responses, we selected transcriptomic datasets with close experimental conditions to ours available on the Gene Expression Omnibus (GEO) repository for comparison **(Figure S11A and B)**. For salt stress, we compared our data to the GSE307070 and GSE339580 datasets for which seedlings were grown on 150mM NaCl for 3h and 200mM for 1h, respectively **(Figure S11A)**. For heat stress, we compared our data to the GSE239833 and GSE339580 datasets which used 37°C and 39°C for 1h stress condition, respectively **(Figure S11B)**^40^. Notably, we observed no correlation between the Log_2_FC from our PUP-IT experiments and the Log_2_FC from transcriptomics, indicating that most of the enrichment observed using PUP-IT corresponds to post-translational responses to stress.

To determine which cellular processes were associated with PI4P and PI(4,5)P_2_ upon the abiotic stresses, we performed a GO term enrichment analysis. As expected, GO terms associated with protein localization and vesicular trafficking were enriched for PI4P and PI(4,5)P_2_ for both salt and heat stress **(Figure 5C and D, S10A and B, Supplementary Dataset 9-10)**. We also confirmed the presence of proteins associated with salt and heat stress **(Figure 5E and F)**. More surprisingly, upon salt stress, PI4P was associated with processes related to cell morphology and cytoskeleton organization while PI(4,5)P_2_ was associated with chloroplast movement and response to stimulus and phosphorylation. However, upon heat stress, response to stimulus and phosphorylation are associated with PI4P only. We also notice an increase in the proportion of transmembrane proteins enriched for PH^FAPP1^ and 2xPH^PLC^ in stress versus control conditions, in particular for PH^FAPP1^ in response to heat. (**Figure S11C and D**). Altogether, our results show that PIP dynamics and their associated proteome are context-dependent. We note that our datasets of PI effectors during salt and heat stress provide a platform to explore lipid-related changes during important stress responses in plants.

## Discussion

Despite the central role of phosphoinositides (PIPs) in membrane organization and signaling, relatively little is known about the proteins that are associated with specific PIP species, particularly in plants. This knowledge gap reflects, in part, the difficulty of systematically identifying lipid-associated proteins *in planta*. Similar questions have been addressed in animal systems using functional lipid probes^41^; however, the application of such probes in plants remains limited. This likely reflects challenges posed by the cell wall, which may restrict probe penetration, as well as uncertainties regarding the extent to which exogenous probes faithfully mimic endogenous lipids. Although cell wall-free protoplast systems can circumvent some of these technical limitations, they do so at the expense of preserving the native cellular context.

To address this challenge, we establish a PUP-IT-based proximity-labeling approach to map PIP-associated proteomes, providing a framework for investigating these interactions in their cellular context. While previous proximity labelling study focused on proteins that enzymatically regulate PIP metabolism^42,43^, we used LBDs to capture broader populations of proteins associated with PI4P and PI(4,5)P₂-containing membrane environments. Encouragingly, our approach revealed distinct sets of proteins linked to each lipid species, including factors involved in signaling, transport, and membrane organization. Notably, transmembrane proteins represented a substantial fraction of the PI4P-associated proteome, suggesting that PI4P-enriched membrane environments may be associated with specific populations of receptors, kinases, and transporters. Whether this observation reflects a functional requirement for PI4P in the activity of these proteins or instead results from the formation of PI4P-rich membrane domains during processes such as endocytosis remains to be determined.

To gain mechanistic insight into how some of the identified lipid-associated proteins engage with PIPs, we combined molecular dynamics simulations with *in vivo* analyses of ARL1 and HIR1. Our results support models in which these proteins use distinct structural features to associate with PIP-containing membranes. For ARL1, simulations and localization experiments point to a positively charged N-terminal surface as the main membrane-binding region, with the C-terminal region potentially contributing to membrane association or orientation. In the case of HIR1, simulations identified positively charged residues at distinct interfaces that may mediate interactions with PIPs. Mutating these residues altered HIR1 plasma membrane sorting and nanodomain organization, supporting a role for PIP interactions in its membrane targeting or stability. However, the persistence of some nanodomain association in the mutated proteins suggests that PIP interactions act alongside other determinants, including HIR1 oligomerization and other membrane components. Indeed, oligomerization status, as well as, membrane sterol content and very-long-chain fatty acid (VLCFA) composition influence HIR2 nanodomain formation^36^. Together, these findings provide some first mechanistic models for how PIP interactions contribute to the membrane association and spatial organization of distinct proteins.

Finally, profiling PIP-associated proteomes under salt and heat stress revealed substantial changes in the proteins associated with PI4P and PI(4,5)P₂. These changes included proteins implicated in vesicular trafficking, lipid signaling, chloroplast movement, and nuclear protein transport, suggesting that PIP-associated protein networks respond to diverse cellular processes during stress. Notably, several previously characterized PIP-dependent stress regulators, including ERD7, FERONIA or PIP5K7 could be re-identified through our approach^44,45^. Moreover, the lack of correlation between protein enrichment and transcriptomic changes under comparable conditions further suggests that these changes cannot be explained by transcriptional responses alone. Nevertheless, the functional significance of the observed associations remains to be established, and our data do not demonstrate that the identified proteins directly mediate stress tolerance. Rather, these datasets provide a starting point for investigating how PIP-associated protein networks change during environmental stress and for identifying candidate effectors involved in abiotic responses.

Although proximity labeling cannot establish direct lipid binding, the approach provides a means of identifying candidate effectors that would be difficult to predict from putative lipid-binding domains, or to capture using conventional biochemical approaches due to weak or transient interactions. The resulting datasets thus offer a resource for investigating the largely unexplored landscape of lipid-associated proteins. More broadly, our work provides a proof of concept for applying proximity-labeling approaches to investigate lipid-associated protein networks across diverse eukaryotic systems.

## Methods

### Growth conditions and plant materials

The Columbia (Col-0) ecotype of Arabidopsis thaliana was used in this study. Arabidopsis thaliana and Nicotiana benthamiana plants were grown in soil in a greenhouse under conditions (16 h light/21 °C and 8 h dark/19 °C). All plant lines used in the study are presented in Table Supp 4.

For transient expression in Nicotiana benthamiana, agrobacterium strain GV3101 carrying the respective destination clones were cultured and co-infiltrated into *N. benthamiana* leaves at OD600 = 0.5 to 0.8 in infiltration buffer.

For in vitro work, Arabidopsis thaliana seeds were surface sterilized before plating on half Murashige and Skoog media containing the appropriate selection. Basta and hygromycin were used at 10mg.L^-1^ and 30mg.L^-1^, respectively. Seedlings were grown in a growth chamber under long-day conditions (16 h light/21 °C and 8 h dark/19 °C) on solid 1/2 Murashige and Skoog (MS) media with 1% sucrose for phenotyping.

### Plasmid construction

All DNA vectors and primers used in this manuscript are presented in Supplementary Table Supp 5 and 6. The constitutive and inducible GFP-PafA lines were described in previous studies^18,20^. The wave marker lines for endomembrane compartments were described in Geldner *et al*., 2009^39^.

To create the PUP-IT entry clones, fragments of MP-mCherry, PH^FAPP1^ E50A, PH^OSBP^, 1xPH^PLC^, 2xPH^PLC^, and TubbyC were amplified from genomic DNA extracted from transgenic lines developed in Simon *et al*., 2014 using Phusion polymerase. 2xPH^PLC^ fragment was amplified as two fragments. PIP5K2 CDS fragment was amplified from Arabidopsis seedling cDNA using Phusion polymerase. All PCR fragments were introduced in a TOPO entry vector containing a Ubiquitin10 promoter, a GS linker, the coding sequence of the PafA and an HSP terminator by Gibson assembly (NEBuilder HiFi DNA assembly mix, New England Biolabs). Subsequently, the ORF was cloned by LR cloning (Gateway LR Clonase II Enzyme mix, Invitrogen) into a destination vector carrying a FLAG::PUP(E) expressed under a constitutive (pMDC32) or a β-estradiol inducible (pFM) 35S promoter.

For reporter lines, HIR1, CPK27, PATL5, BIR2, GPA1, ARG1, and ARL1 CDS fragments were amplified from Arabidopsis seedling cDNA using Phusion polymerase and introduce into a pGGC000 entry vector using GreenGate technology or Gibson assembly. When necessary silent mutations were introduced to remove *BsaI* sites by directed mutagenesis using the indicated primers (Table 9). For ARL1 truncated reporter lines, corresponding CDS fragments were amplified from ARL1/pGGC and reintroduced into pGGC000 entry vector.

Point mutagenesis on HIR1 was performed by amplifying HIR1/pGGC with primers containing the corresponding point mutations, followed by *dpn1* digestion before E. coli transformation (Table 11). Full constructs were assembled into destination vector pZ001 using GreenGate technology and the entry vectors pGGA004 (*35S (Cauliflower mosaic virus 35S*) promoter), pGGA006 (*UBQ10* (*UBIQUITIN10*) promoter), pGGB003 (*B-dummy*), pGGD002 (*D-dummy),* pGGE009 (UBQ10 terminator), pGGF002 (p35S BastaR:t35S), PGGF005 (pUBQ10:Hygro:tOCS)^46^.

Finally, destination vectors were transformed into the Agrobacterium strain GV3101 together with pSoup helper plasmid for plant transformation by floral dipping.

### Protein Extraction and Enrichment

For samples in *N. benthamiana*, the 2 to 3g of tissues were collected 2 days after infiltration and ground to powder in liquid nitrogen. The powder was resuspended in extraction buffer (1:3 [W/V], [20 mM Tris-HCl, pH 7.4, 150 mM NaCl, 5 mM MgCl2, 1 mM EDTA, 1% Triton X-100, and 0.1% protease inhibitor cocktail (Roche)] and incubated for 30 min on ice, and centrifuged at 4 °C at 16000g for 30 min. The supernatant was divided into three technical replicates for the rest of the procedure. Each sample was incubated with 100 µL equilibrated anti-Flag antibody coupled magnetic beads (Thermo Scientific, A36797) for 2 h at 4 °C. Beads were washed three times with extraction buffer, three times with wash buffer [20 mM Tris-HCl, pH 7.4, 150 mM NaCl, 5 mM MgCl2, 1 mM EDTA, and 0.1% protease inhibitor cocktail (Roche)] and three times with 1 X PBS and stored at –70 °C until LC-MS/MS analysis.

For Arabidopsis constitutive samples, 1 to 2 g of Arabidopsis seedlings from two independent transgenic lines were used. Similarly to *N. benthamiana* samples, tissue was grounded, resuspended with extraction buffer (1:3 [W/V]), [50 mM HEPES pH 7.5, 150 mM NaCl, 10 mM NaF, 5 mM EDTA, 1% Triton X-100, 1% PVP40, 0.1% protease inhibitor cocktail (Roche)] and incubated for 30 min on ice before centrifugation at 16 000 g at 4 °C for 30 min and incubation with 100 µL equilibrated anti-Flag antibody coupled magnetic beads.

Three independent pFM inducible transgenic seedlings were induced by adding 10 µM β-estradiol in 10mL liquid half-MS onto 8-days old seedlings. After 6 h, 10 mL of 400 mM NaCl liquid half-MS was added for salt stress while 10mL liquid half-MS was added to the rest of the plates. Finally, plates were moved to an incubator at 35°C for heat stress. After 2 h, tissue was collected, frozen in liquid nitrogen and ground. 1 ml of ground powder per sample was extracted with 1.5 ml extraction buffer, incubated for 30 min on ice before centrifugation at 16 000 g at 4 °C for 30 min and incubation with 25 µL equilibrated anti-Flag antibody coupled magnetic beads.

Beads were washed three times with extraction buffer, then three times with wash buffer [50 mM HEPES pH 7.5, 150 mM NaCl, 10 mM NaF, 5mM EDTA, 0.1% protease inhibitor cocktail (Roche)] and three times with 1 X PBS buffer.

To assess pupylation labelling, 10µl aliquot per samples were taken during the last wash and analyzed by western blot with FLAG antibody (Merck, F3165, diluted 1:2000).

### Protein Digestion and Mass Spectrometry

Washed beads were first eluted for 30 min at 37°C with 2 M Urea, 50 mM Tris-HCl pH 7.5, 2 mM DTT, 20 µg mL^−1^ trypsin. The eluates were mixed with an equal in 2 M Urea, 50 mM Tris-HCl pH 7.5, 10 mM Chloroacetamide for overnight incubation. Samples were acidified to 1% TFA and loaded on Evotips (Evosep). Peptides were separated on 15 cm, 150 µm ID columns packed with C18 beads (1.5 µm) (Pepsep) on an Evosep ONE HPLC applying the “30 samples per day” method while the column temperature was maintained at 50°C and injected via a CaptiveSpray source and 20 µm emitter into a timsTOF pro2 mass spectrometer (Bruker) ran in PASEF mode. MS data were collected over a 100-1700 m/z range with a TIMS mobility range of 0.6-1.6 1/K_0_. Ion mobility was calibrated using three Agilent ESI-L Tuning Mix ions: 622.0289, 922.0097, and 1221.9906. TIMS ramp and accumulation times were set to 100 ms each, with 10 PASEF ramps recorded for a total cycle time of 1.17 s. MS/MS target intensity and intensity threshold were set to 20.000 and 2.500, respectively. An exclusion list of 0.4 min for precursors within 0.015 m/z and 0.015 V cm^−2^ width was activated.

### Peptide search

Raw timsTOF-PASEF mass spectrometry data were processed using FragPipe (v23.1)^47^. MS/MS spectra were searched with MSFragger against the Uniprot *A. thaliana* reference proteome or the *N. benthamiana* reference proteome from Kourelis et al., 2019 (with added contaminants and reversed decoys), allowing up to three missed tryptic cleavages and at least one tryptic terminus (i.e. semi-tryptic search)^48^. Precursor and fragment mass tolerances were set to 20 ppm, with isotope errors of 0/1/2; mass calibration and parameter optimization was enabled. Carbamidomethylation of cysteine (+57.02146 Da) was set as fixed modification; variable modifications were methionine oxidation, protein N-terminal acetylation and the pupylation remnant GGE on lysines (+243.08553 Da). Spectra were rescored with MSBooster^49^. PSMs were validated with Percolator and protein inference was performed with ProteinProphet; results were filtered at 1% PSM, peptide, and protein FDR using Philosopher. Label-free quantification was performed with IonQuant using MaxLFQ normalization, with match-between-runs enabled; peptide and protein quantification were filtered at 1% FDR.

### Differential enrichment analysis

Proteomics data analysis was performed in R (v4.5). Contaminant and decoy sequences were removed; the remaining peptide intensities were log-transformed and median-centered. Proteins identified by only one peptide were filtered out. Missing values were imputed on peptide level using the PIRAT package (v1.9.0), which determines an empiric instrument detection level and takes into account all peptide intensities for each protein, efficiently imputing both missing-at-random and missing-not-at-random peptides^50^. Since PIRAT imputation performs poorly when all peptides of a protein are completely absent from a whole sample group, such proteins were imputed with a MinProb approach, i.e. a value sampled from a normal distribution with a mean of the experiment-wide 0.5% quantile and a standard-deviation of 0.05. Differential enrichment between groups was calculated directly from peptide levels using MSqRob (v1.20.0), including random effects for the sample, peptide sequence and experiment (where applicable)^51^. Protein intensities for visualisation were estimated using robust summarisation; where protein levels needed to be compared across experiments, batch effects were removed using the the R package ‘limma’ (v3.68.4)^52,53^. Arabidopsis homologs of *N. benthamiana* proteins were identified using MMseqs2^54^. The code for the entire processing pipeline is available under https://doi.org/10.5281/zenodo.24013703.

### GO term enrichment analysis

Functional enrichment analysis was performed using the STRING database version 12.0 to identify significantly overrepresented Gene Ontology (GO) terms^55^. The GO term network was visualized using ggraph extension of ggplot2.

### Pharmacological treatments

For PAO treatment, 5 to 7-days old seedlings were incubated in liquid half-MS with 60 µM PAO (Sigma-Aldrich, P3075, stock solution 60 mM in DMSO) or the equivalent volume of DMSO for 30 min or 60 min before observation.

For BFA treatment, seedlings were placed in liquid 1/2 MS and treated with 50 µM BFA (Sigma-Aldrich, B7651, stock solution 5 mg.mL^−1^ in DMSO) or the equivalent volume of DMSO for 60min before observation.

### Artificial microRNA induction

5-days old seedlings were transferred on half-MS media supplemented with 1% sucrose and 5 µM β-estradiol (Sigma-Aldrich, P3075, stock solution 60 mM in DMSO) or the equivalent volume of DMSO and let to grow for 2 days before observation. Intracellular to plasma membrane ratio were calculated by the mean grey value along the basal plasma membrane on the mean grey value of the intracellular space.

### Microscopy settings and image analysis

Microscopy images were taken using Confocal Microscope 3i CSU-W SoRa Spinning Disk (Camera: Andor iXon Life 888 1024×1024 EMCCD) and processed by Fiji for image analyses. mNEON and mScarlet were excited by laser 488 nm (20%, 400 ms) and laser 561 nm (20%, 400 ms), respectively and observed under a 60X or 100X oil lens. PM/intracellular ratio were calculated by the mean grey value along the basal PM on the mean grey value of the intracellular space.

For co-localization with the wave markers, images were taken using simultaneous imaging with mNEON and mCherry excited by laser 488 nm (20%, 400 ms) and laser 561 nm (20%, 400 ms), respectively and observed under a 60X oil lens.

TIRF images were taken on a Zeiss Elyra PS.1. mNEON was visualized using laser 488 nm, BP 495-575, LP 750, 40ms exposition time, gain 8, under a 100X oil lens. mScarlet3 was visualized using laser 561 nm, BP 570-650, LP 750, 250ms exposition time, gain 30 under a 100X oil lens. CV was calculated by dividing the standard deviation by the mean of the signal intensity along 10µm linear profiles.

### Structure prediction

Structures of oligomeric assemblies were predicted using AlphaFold3^56^. The quality and local confidence of the obtained models were assayed based on the pLDDT score, while the global confidence and interaction between subunits were assessed by the ipTM and predicted alignment error (PAE) metrics.

### Coarse-Grained Molecular dynamics simulations and trajectory analysis

All simulations were performed using GROMACS version 2023.3.

Coarse-grained (CG) molecular dynamics simulations were performed using the MARTINI 2.2 force field. For the HIR1 14-mer, the initial atomistic structure of the protein was obtained using the AF3, oriented towards a membrane using the PPM 3.0 webserver^57^ and converted to a coarse-grained representation using the Eldenyn22 version of the Martini 2.2 forcefield via the CharmmGUI server. Posttranslational palmitoyl and myristoyl were added to each subunit using the martinize script developed by Atsmon-Raz *et al*., 2017^58^. A membrane containing 37% POPC, 37% POPE, 10% POPS, 10% POPA, 5% POP1 and 1% POP2 in the cytosolic leaflet and equal amount of POPC and POPE in the extracellular leaflet was generated using the INSANE tool. The protein complex together with the membrane was solvated with coarse-grained water beads. Sodium and chloride ions were added to achieve electroneutrality and a salt concentration of 0.15 M.

For the ARL1 dimer, the initial atomistic structure of the protein was obtained using AF3, truncated to conserve the amino acids 1 to 354, oriented using the PPM 3.0 webserver and converted to a coarse-grained representation using the Eldenyn22 version of the Martini 2.2 forcefield via the CharmmGUI server together with a lipid bilayer consisting of 37% POPC, 37% POPE, 10% POPS, 10% POPA, 5% POP1 and 1% POP2 in each leaflet^57,59^. The protein complex together with the membrane was solvated with coarse-grained water beads. Sodium and chloride ions were added to achieve electroneutrality and a salt concentration of 0.15 M.

Prior to equilibration, all systems were subjected to energy minimization using the steepest-descent algorithm implemented in GROMACS. Minimization was performed for a maximum of 5,000 steps using periodic boundary conditions in all directions.

Equilibration was conducted in the NPT ensemble with position restraints applied to the protein and lipid headgroups.

Production simulations were performed for 10 μs using a time step of 20 fs. Temperature was maintained at 303 K using the velocity-rescale thermostat with a coupling constant of 1.0 ps. Pressure was maintained at 1 bar using the Parrinello-Rahman barostat with semi-isotropic coupling and a relaxation time of 12 ps.

Lipid density analyses were performed using the gromacs in built gmx density function. Protein-lipid contacts were calculated using the Python package PyLipID with a cutoff distance of 0.7 nm^60^.

### Atomistic Molecular dynamics simulations and trajectory analysis

The atomistic structure of the protein was obtained using AF3 and truncated to conserve the indicated protein domain. Atomistic membrane systems were constructed using the CHARMM-GUI Membrane Builder^59,61^. Proteins were described using the CHARMM36m force field, while lipids were parameterized using the CHARMM36 lipid force field.

The protein was positioned above a POPC bilayer containing 10% POPI14 (PI4P) and solvated with TIP3P water molecules. Potassium and chloride ions were added to neutralize the system and produce a final ion concentration of 0.15 M.

Energy minimization was performed using the steepest-descent algorithm for 5,000 steps. The system was then equilibrated through a six-step protocol involving gradually decreasing positional restraints on protein and lipid atoms.

Production MD simulations were performed using GROMACS version 2023.3. Simulations were conducted in the NPT ensemble using a 2fs integration time step. Temperature was maintained at 303 K using a velocity-rescaling thermostat, while pressure was controlled at 1 atm using the Parrinello-Rahman barostat with semi-isotropic pressure coupling. Pressure was maintained at 1 bar using the stochastic cell-rescaling (C-rescale) barostat with semi-isotropic pressure coupling.

Each system was simulated for 500 ns, and three independent replicas were generated using different initial velocity distributions.

Lipid density analyses were performed using the gromacs in built gmx density function. Protein-lipid contacts were calculated using the Python package PyLipID with a cutoff distance of 0.7 nm^60^.

### Differential Gene Expression Analysis

Transcriptomic datasets were downloaded from Gene Expression Omnibus (GEO) repository. When only raw gene counts were available, differential expression analysis was carried out on count data from three control samples and three treatment samples. Differential gene expression analysis was performed using the DESeq2 package (version 1.52.0). Normalization of sequencing depth and estimation of gene-wise dispersion parameters were carried out using the default DESeq2 workflow. Negative binomial generalized linear models were fitted for each gene, and differential expression between treatment and control samples was assessed using the Wald test. Genes were considered significantly upregulated in the treatment group when they satisfied Log_2_FCf > 1and adjusted p-value (FDR) < 0.05.

## Acknowledgements

L.C.N. acknowledges EMBO postdoctoral fellowship (ALTF 629-2021) and Novo Nordisk postdoctoral grant (NNF24OC0094901). S.P. acknowledges Villum Investigator (Project ID: 25915), Novo Nordisk Laureate (NNF19OC0056076), Novo Nordisk Data Science (NNF0068884) grants. L.B. acknowledges EMBO postdoctoral fellowship (ALTF 37-2022) and support through the Wallenberg Initiatives in Forest Research (WIFORCE; KAW 2020.0240). We acknowledge the Danish e-infrastructure Consortium (DeiC) for awarding this project access to the LUMI supercomputer, owned by the EuroHPC Joint Undertaking, hosted by CSC (Finland) and the LUMI consortium through DeiC. Mass spectrometry-based proteomic analyses were performed by the Proteomics Research Infrastructure (PRI) at the University of Copenhagen (UCPH), supported by the Novo Nordisk Foundation (NNF) (grant agreement number NNF19SA0059305. We thank Dr. Alexis Lebecq for discussion and comments on the manuscript.

## Author contributions

L.C.N conceived, designed and performed experiments, analyzed data and wrote the manuscript. P.V. performed experiments and helped generate transgenic lines. M.N and R.P provided technical support for the molecular dynamics. L.B performed the mass spec analyses. S.P. provided guidance and suggestions for the study. All authors have edited and approved the manuscript.

## Materials & Correspondence

Plasmids and seeds are available directly from Lise Noack upon request.

## Data availability

The Raw proteomics data generated in this study have been deposited in the server. The scripts for the complete proteomics analyses in study are available under https://doi.org/10.5281/zenodo.24013703.

## Supplementary Information

**Figure S1.**
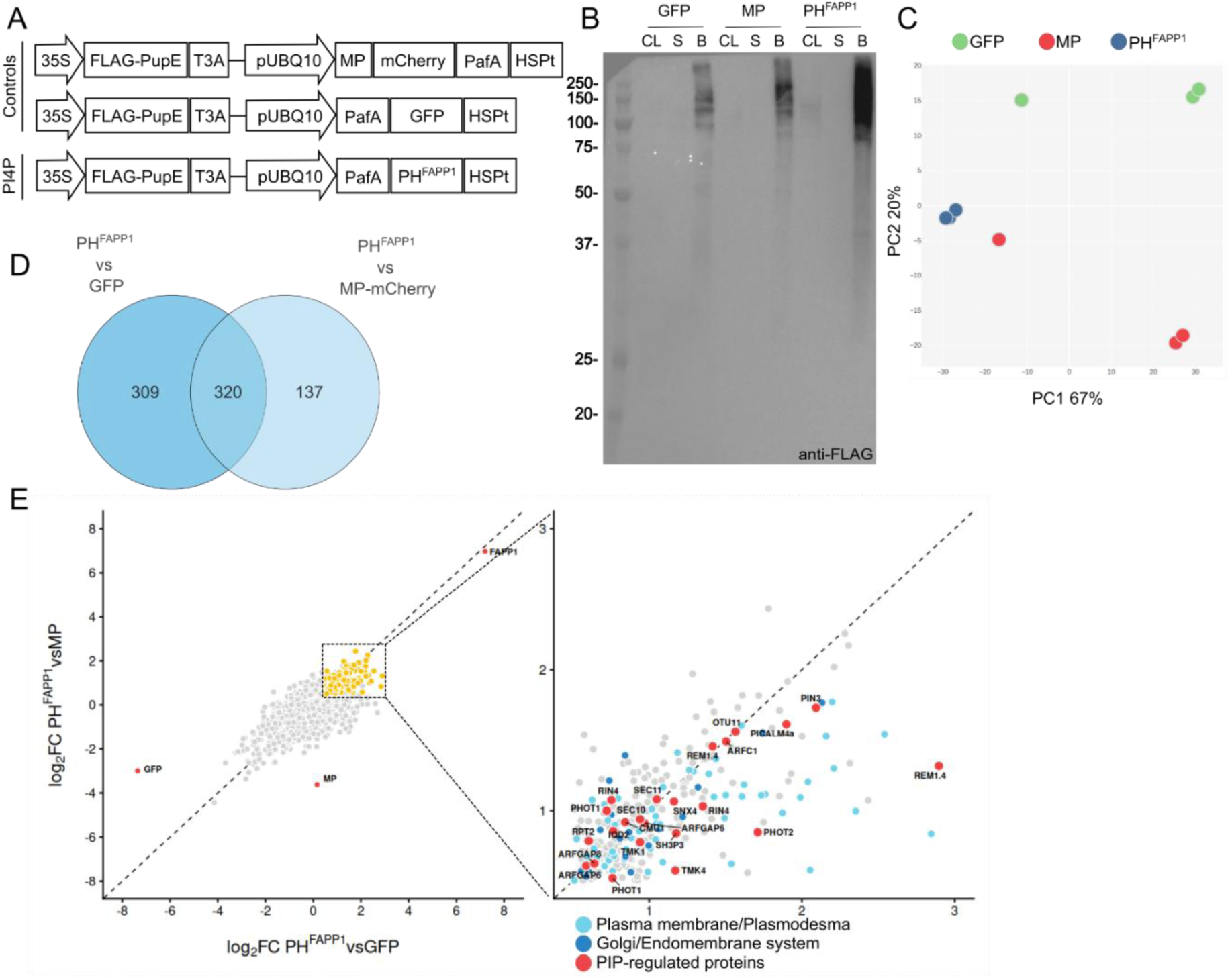
Identification of the PI4P-associated proteome by Pupylation-Based Proximity Labeling in N. benthamiana. **A.** Schematic representation of the PUP-IT constructs used in N. benthamiana. For each construct the FLAG-PupE expression is driven by a 35S promoter and terminated by a T3A terminator while the bait expression is driven by a UBQ10 promoter and terminated by an HSP terminator. **B.** Anti-FLAG western blot of PUP-IT constructs transiently expressed in Nicotiana benthamiana leaves. CL, Crude lysate; S, supernatant; B, beads. **C.** Principal component analysis on normalized protein intensity of PUP-IT constructs transiently expressed in Nicotiana benthamiana leaves. **D.** Venn Diagram of the proteins found with a log_2_FC>0.5 and an adjusted p-value<0.05 in the different comparisons. **E.** Volcano plots showing proteins significantly enriched in PH^FAPP1^ versus controls in yellow. Baits are shown in red. In the close-up, proteins which have already been associated with PI4P in scientific literature are indicated in red. Proteins with GO term associated with the PM or plasmodesmata and with the Golgi or endomembrane (*trans*-Golgi Network, endosomes) are shown in light blue and dark blue, respectively.

**Figure S2.**
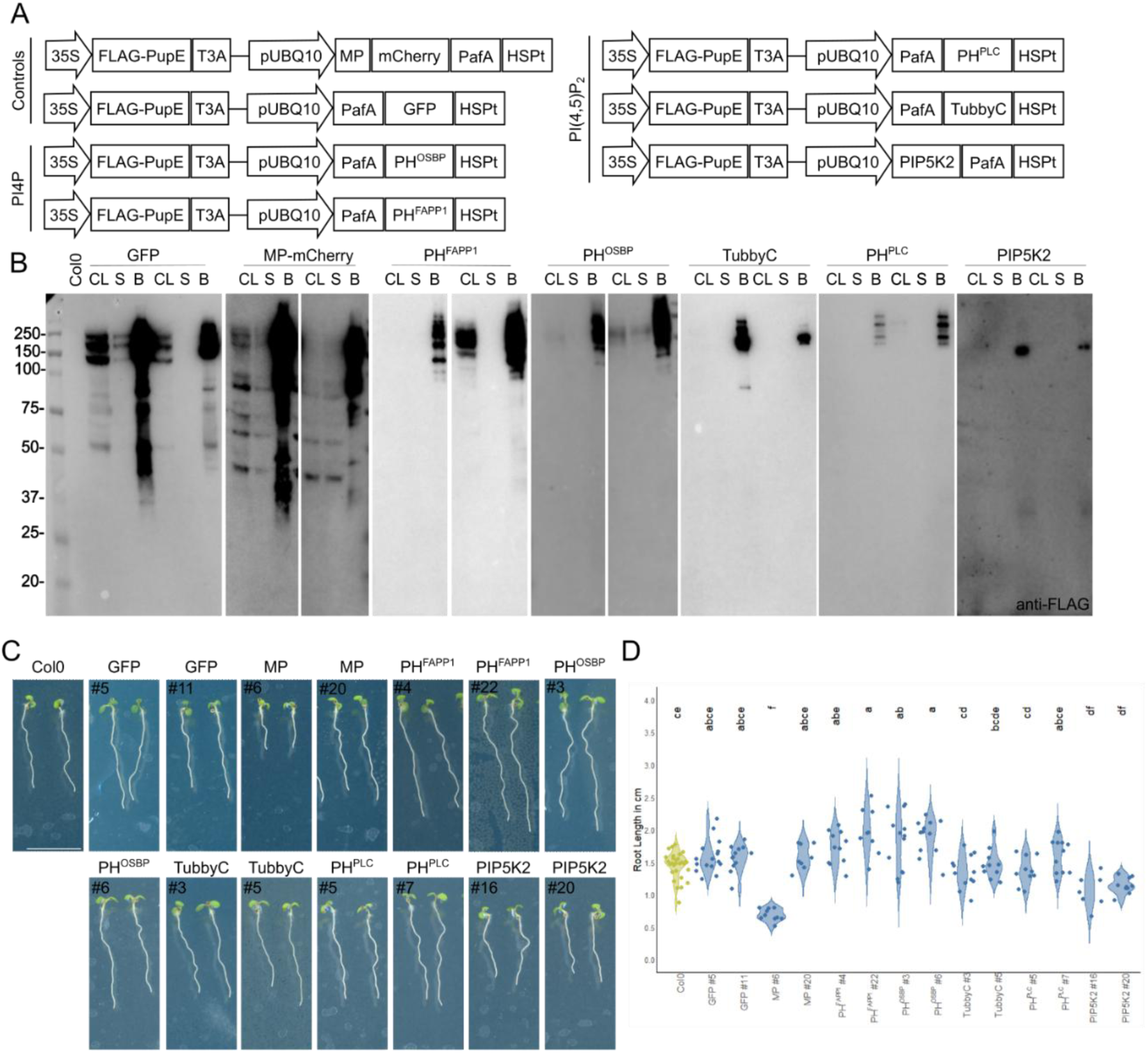
Validation of the PUP-IT constructs used to identify the PIP-associated proteomes in A. thaliana. **A.** Schematic representation of the PUP-IT constructs used in *A. thaliana*. For each construct the FLAG-PupE expression is driven by a 35S promoter and terminated by a T3A terminator while the bait expression is driven by a UBQ10 promoter and terminated by an HSP terminator. **B.** Anti-FLAG western blot of PUP-IT constructs expressed in *A. thaliana* seedlings. Two independent lines showing enrichment in the bead fraction compared to supernatant have been selected for each construct. CL, Crude lysate; S, supernatant; B, beads. **C.** 7-day-old *Arabidopsis* seedlings expressing the PUP-IT construct constitutively. Scale bar= 1 cm **D.** Root length measurements of 7-day-old *Arabidopsis* seedlings expressing the PUP-IT construct constitutively. Statistical analysis was done using the Kruskal-Wallis test, followed by Dunn’s test with Benjamini-Hochberg correction.

**Figure S3.**
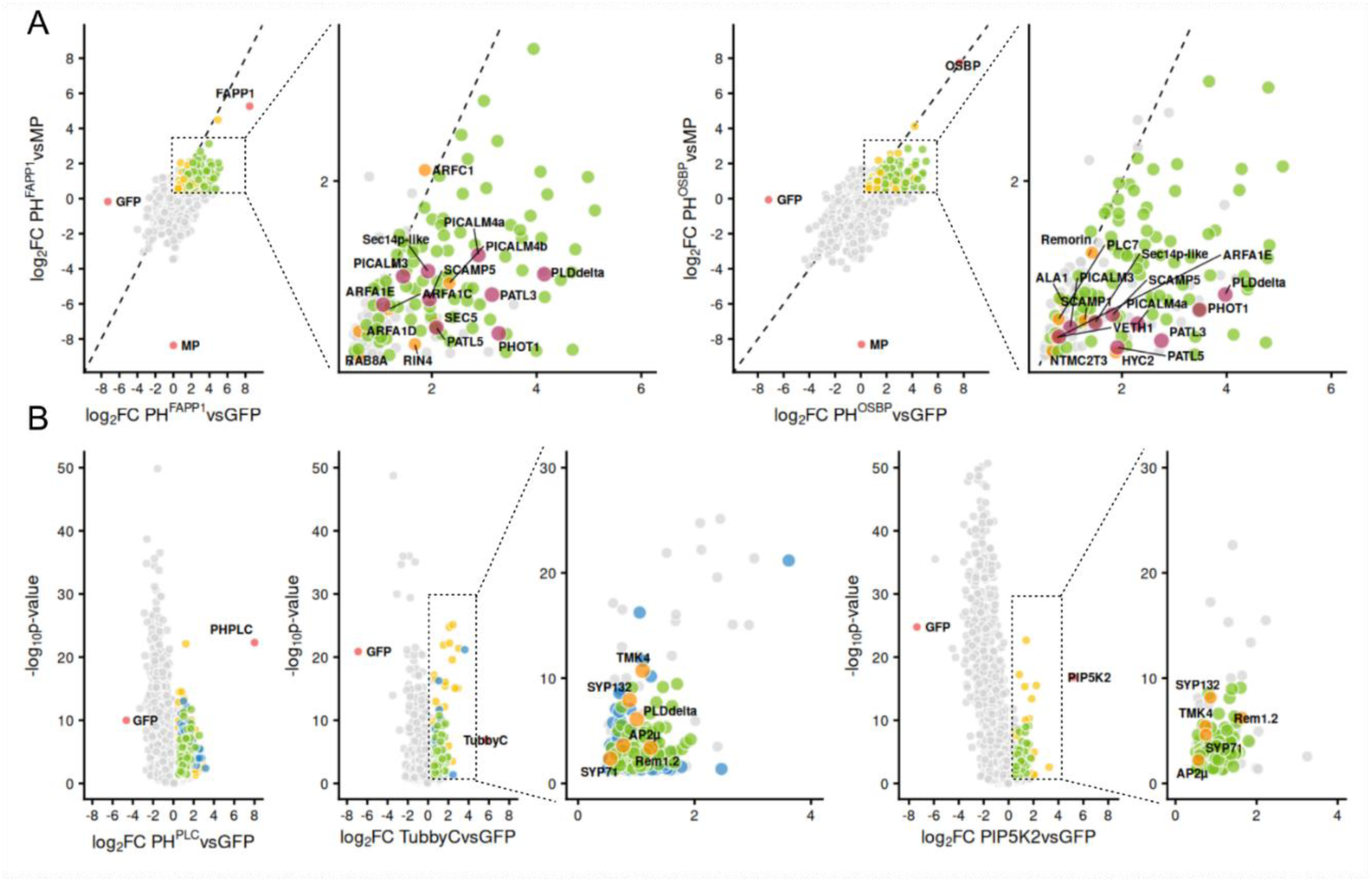
PIP-related proteins enriched in PI4P- and PI(4,5)P_2_-interactomes in A. thaliana. **A.** Volcano plots showing proteins significantly enriched in PH^FAPP1^ and PH^OSBP^ versus controls in yellow. Proteins enriched in both PH^FAPP1^ and PH^OSBP^ are shown in green. Baits are shown in red. In the close-up, proteins which have already been associated with PIPs in scientific literature are indicated in orange when found specifically in one bait and in purple when enriched in PH^FAPP1^ and PH^OSBP^. **B.** Volcano plots showing proteins significantly enriched in PH^PLC^, TubbyC and PIP5K2 versus controls in yellow. Proteins enriched in both PH^PLC^ and TubbyC are shown in blue, while proteins enriched in PH^PLC^, TubbyC and PIP5K2 are shown in green. Baits are shown in red. In the close-up, proteins which have already been associated with PIPs in scientific literature are indicated in orange.

**Figure S4.**
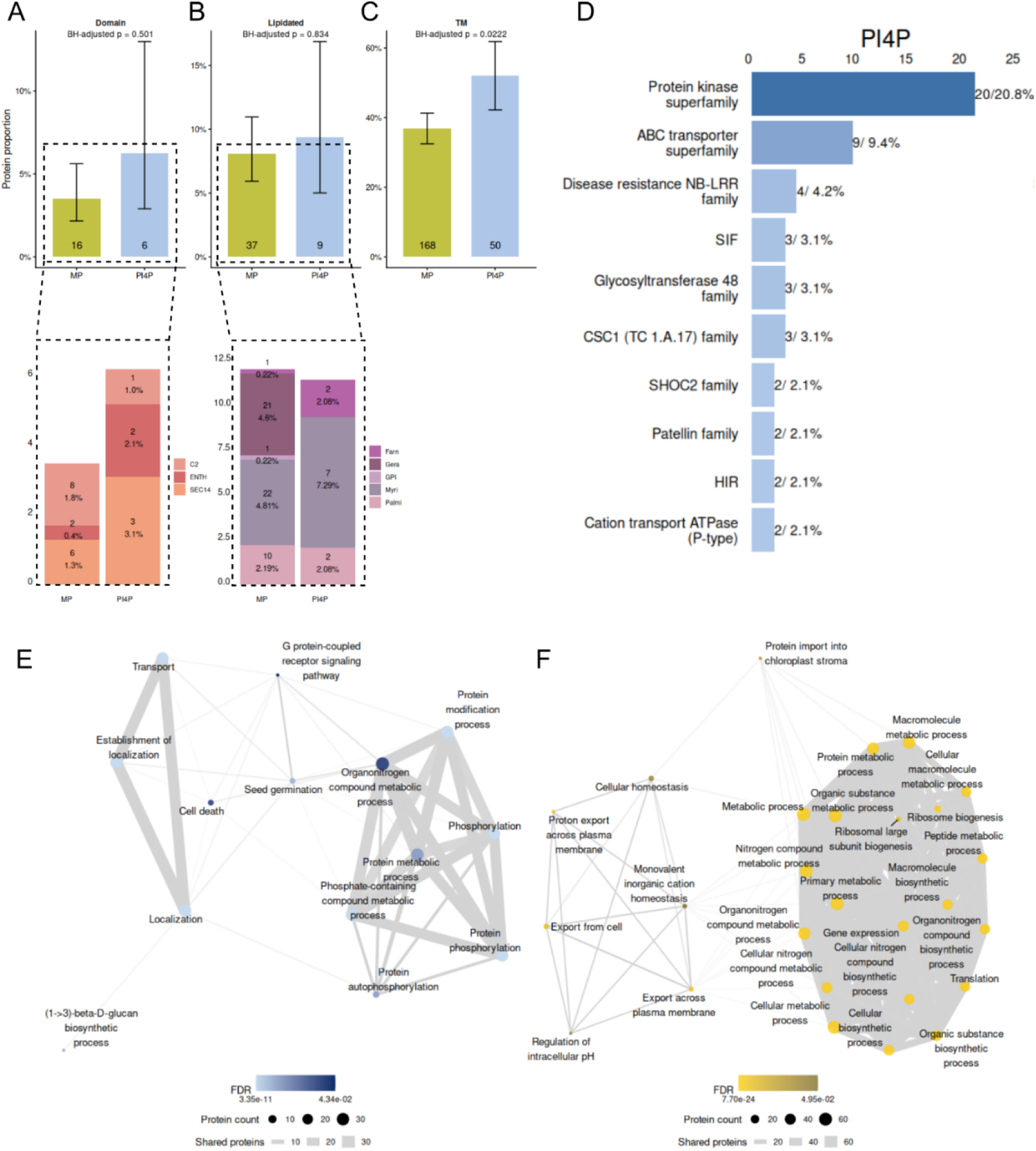
Analyses of proteins enriched in PI4P- and PI(4,5)P_2_-interactomes in A. thaliana A-C. Proportion of proteins containing a lipid binding domain (A), a lipidation site (B) and a transmembrane domain (C) enriched in MP-mCherry (MP) and PI4P-associated proteome. The number of proteins is indicated in each bar. The details of the lipid binding domains (A) and lipidation sites with the proportion and number of proteins are shown below each graph. Farn, Farnelysation; Gera, Geranylgeranylation; GPI, Glycosylphosphatidylinositol anchor; Myri, Myristoylation; Palmi; Palmitoylation. For statistical analysis, Pearson’s Chi-squared test when sample size was above 5 and Fisher’s exact test when sample number was under or equal to 5 were used. P-values were adjusted using Benjamini-Hochberg (BH) method. **D.** Top 10 protein families enriched in PI4P-associated proteome. The proportion and number of proteins are indicated above each bar. **E,F.** Gene Ontology analysis of cellular processes for proteins enriched in the PI4P-(E) and PI(4,5)P_2_-associated (F) proteomes.

**Figure S5.**
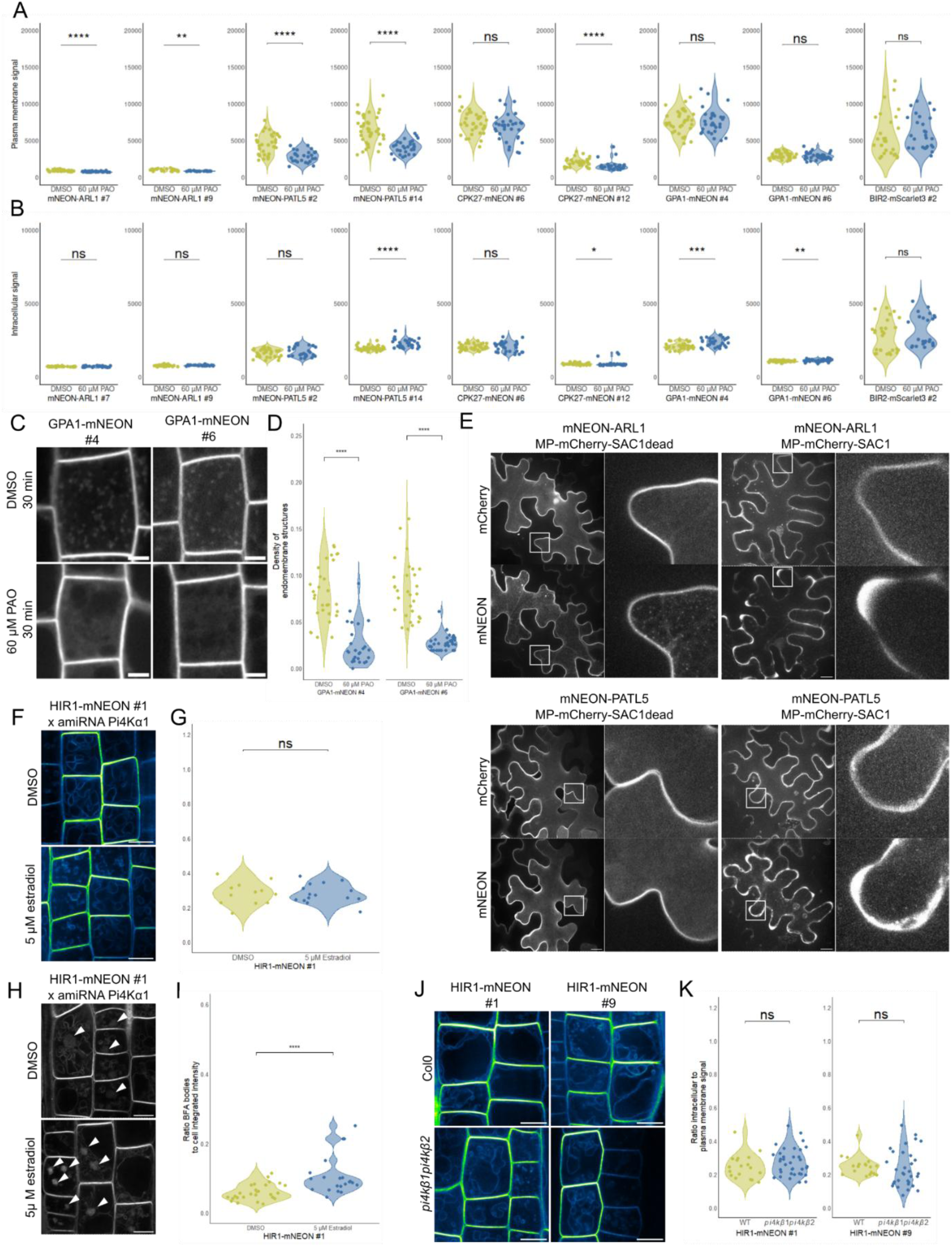
PI4P-associated protein candidates change their distribution patterns and dynamics upon PIP depletion A,B. Plasma membrane signal (A) and intracellular signal (B) of Arabidopsis root epidermal cells expressing pUBQ10:mNeon-ARL1, pUBQ10:mNeon-PATL5, pUBQ10:CPK27-mNeon, pUBQ10:GPA1-mNeon, and pUBQ10:BIR2-mScarlet3 treated with 60 µM PAO for 30 min or the equivalent volume of DMSO. Student’s T test was used to assess statistical significance when normal distribution was confirmed using the Shapiro test. Wilcoxon rank-sum tests were used otherwise (n ≥ 24). **C,D.** Confocal microscopy images (C) and quantification of the density of endomembrane structure (per µm^-2^) (D) of Arabidopsis root epidermal cells expressing pUBQ10:GPA1-mNeon treated with 60 µM PAO for 30 min or the equivalent volume of DMSO. Scale bar = 5 µm. Statistical significance is based on a pairwise Wilcoxon test (n ≥ 23). **E.** Confocal microscopy images of Nicotiana leaf epidermal cells expressing mNEON-ARL1 or mNEON-PATL5 with MP-mCherry-SAC1 or MP-mCherry-SAC1dead. Scale bar = 20 µm **F,G.** Confocal microscopy images (F) and calculated ratio of intracellular to plasma membrane fluorescent signal (G) of Arabidopsis root epidermal cells expressing p35S:HIR1-mNeon carrying the amiRNA_PI4Kα1 treated with 5 µM β-estradiol or the equivalent volume of DMSO. Scale bar = 10 µm. Student’s T test was used to assess statistical significance after normal distribution was confirmed using the Shapiro test. (n ≥ 12). **H,I.** Confocal microscopy images (H) and calculated ratio of intracellular to plasma membrane fluorescent signal (I) of Arabidopsis root epidermal cells expressing p35S:HIR1-mNEON carrying the amiRNA_PI4Kα1 treated with 5 µM β-estradiol or the equivalent volume of DMSO before being treated with 50 µM BFA for 60 min or the equivalent volume of DMSO. Scale bar = 10 µm. Statistical analysis was done using a pairwise comparisons Wilcoxon rank-sum test. (n ≥ 21) **J,K.** Confocal microscopy images (J) and calculated ratio of intracellular to plasma membrane fluorescent signal (K) of Arabidopsis root epidermal cells expressing p35S:HIR1-mNeon in wild-type (Col0) or *pi4kβ1pi4kβ2* genetic background. Scale bar = 10 µm. Student’s T test was used to assess statistical significance when normal distribution was confirmed using the Shapiro test. Wilcoxon rank-sum tests were used otherwise. (n ≥ 18).

**Figure S6.**
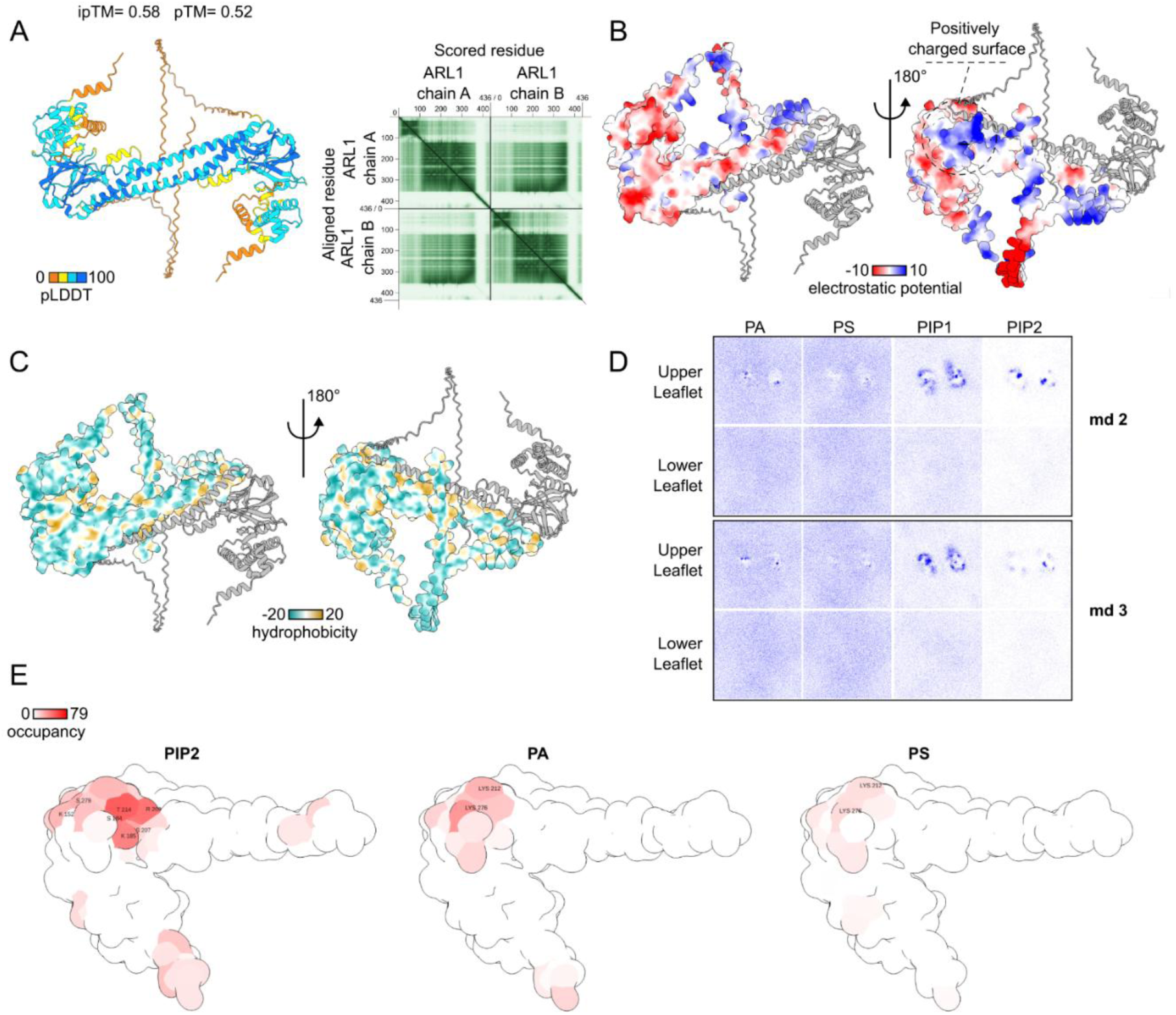
The ARL1 dimer shows lower affinity for PA, PS and PI(4,5)P_2_ than PI4P. **A.** AF3 model of the ARL1 dimer. Local model confidence is shown by the pLDDT score. The global model quality is validated by a low predicted alignment error (PAE). **B.** Surface representation of the ARL1 dimer, colored by the Coulombic electrostatic potential. **C.** Surface representation of the ARL1 dimer, colored according to the molecular lipophilicity potential. **D.** Two-dimensional density (over 10 µs) of PIP1, PIP2, PA and PS in the upper leaflet (adjacent to the protein) and lower leaflet for replicates 2 and 3. **E.** Occupancy of PA and PS mapped onto ARL1(1-354) structural representation. The occupancy is defined as the percentage of frame with contact (distance under 0.7 Å).

**Figure S7.**
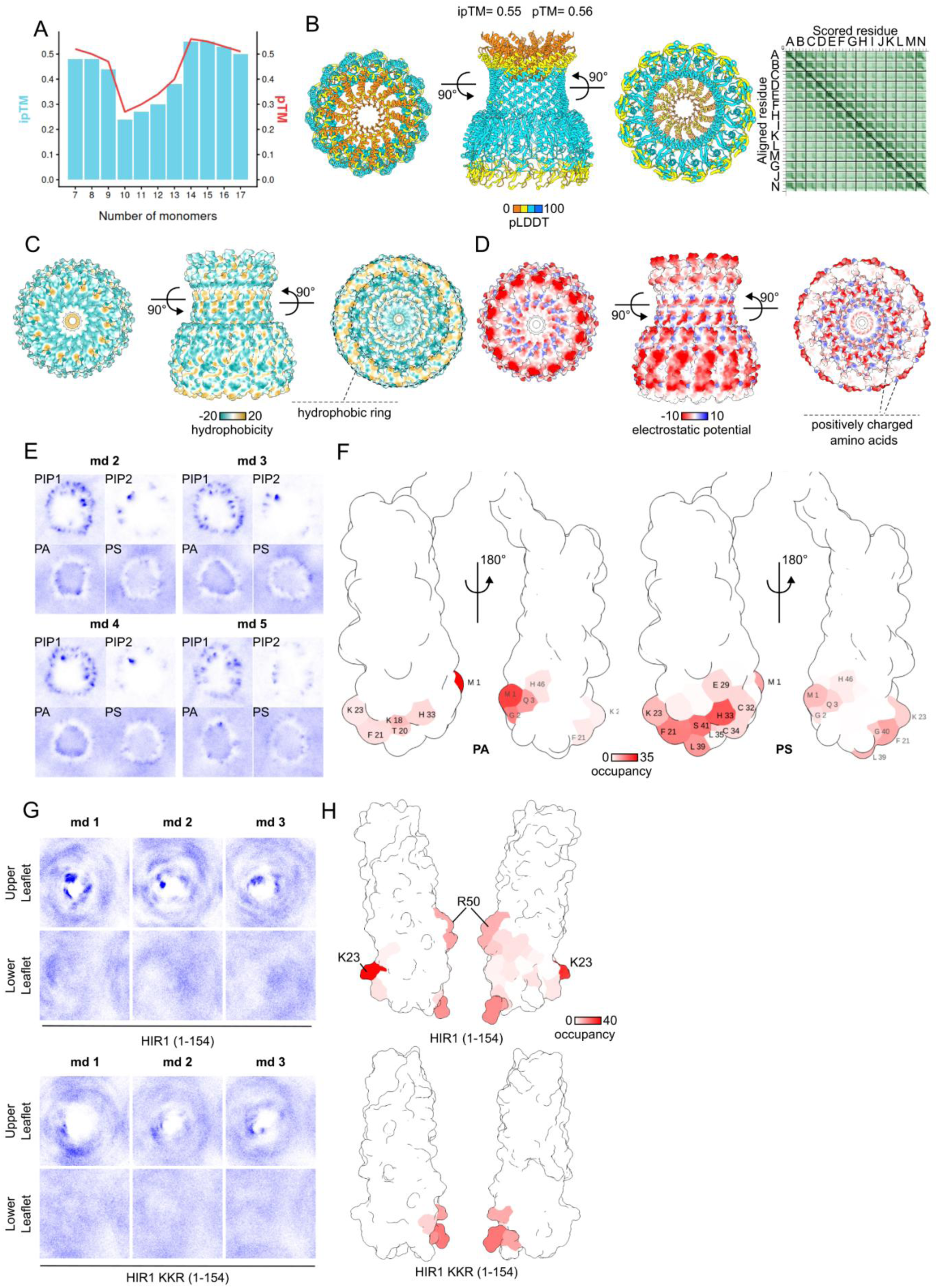
HIR1 oligomer binds PIPs via positively charged amino acids. **A.** AF3 predictions of HIR1 complex stoichiometries containing 7 to 17 subunits of HIR1. **B.** AF2 model of the HIR1 complex containing 14 subunits. Local model confidence is shown by the pLDDT score. The global model quality is validated by a low predicted alignment error (PAE). **C.** Surface representation of the HIR1 14-mer, colored according to the molecular lipophilicity potential. **D.** Surface representation of the HIR1 14-mer, colored by the Coulombic electrostatic potential. **E.** Two-dimensional density (over 10 µs) of PIP1, PIP2, PA and PS in the cytosolic leaflet adjacent to the HIR1 complex for replicate 2 to 5. **F.** Occupancy map of PA and PS on the HIR1 SPFH domain. The occupancy is defined as the % of frame with contact (distance under 0.7 Å). **G.** Two-dimensional density over 500ns of PI4P in the upper leaflet (adjacent to the HIR SPFH domain) and lower leaflet from atomistic MD simulations of the HIR1 SPFH domain (1-154) and the HIR1 KKR SPFH domain (1-154) with a POPC membrane containing 10% PI4P in each leaflet. **E.** Occupancy of PI4P HIR SPFH domain and HIR1 KKR SPFH domain (1-154).

**Figure S8.**
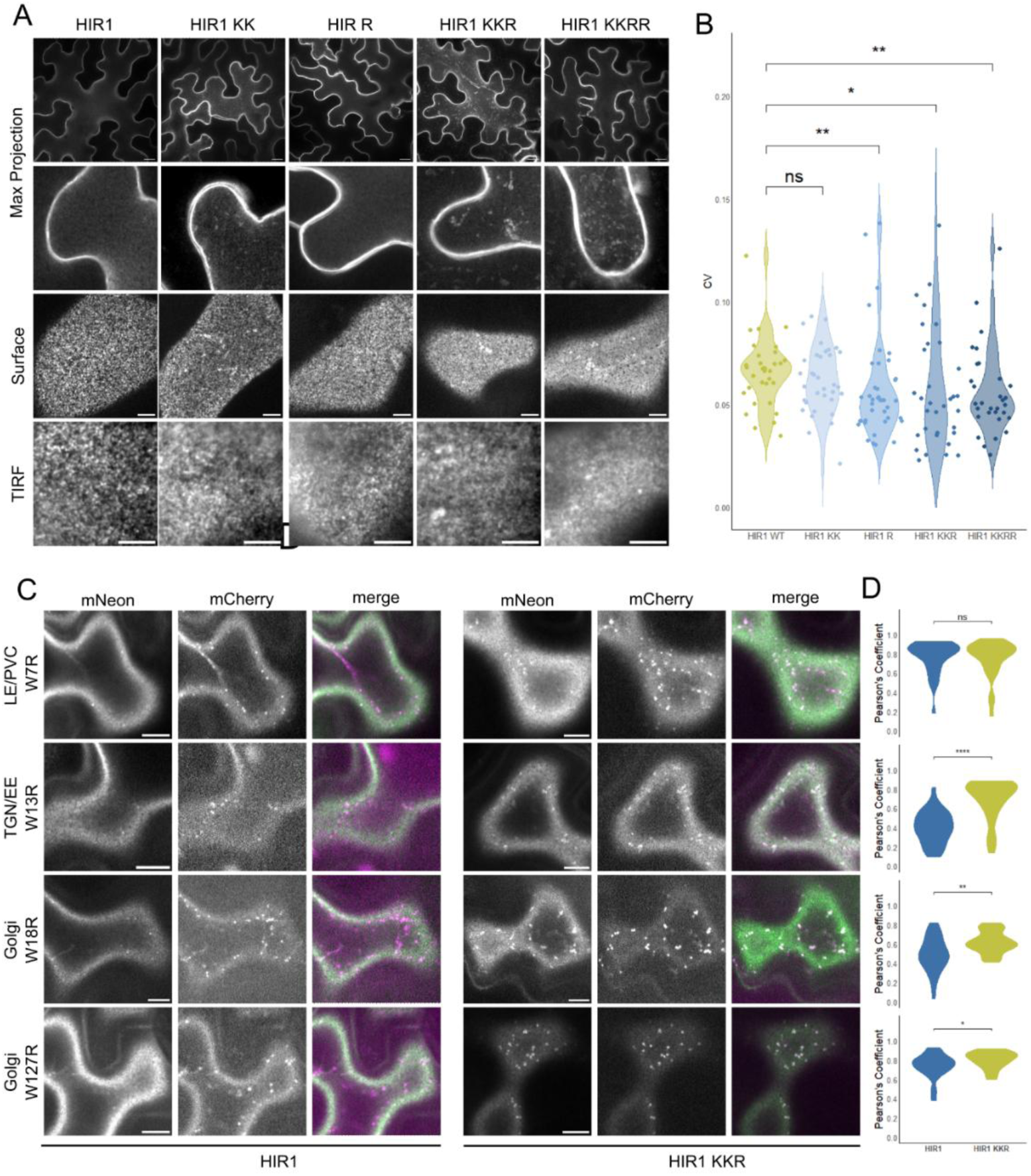
HIR1 positively charged amino acids are necessary for HIR1 stability at the PM in N. benthamiana. **A.** Max projection (top panels), and surface view (middle panel) of confocal microscopy images and TIRF microscopy images (lower panel) of Nicotiana leaf epidermal cells transiently expressing p35S:HIR1-mNEON, p35S:HIR1(K18Q;K23Q)-mNeon (HIR1 KK), p35S:HIR1(R50Q)-mNeon (HIR1 R), p35S:HIR1(K18Q;K23Q;R50Q)-mNeon (HIR1 KKR), p35S:HIR1(K18Q;K23Q;R50Q;R94Q)-mNeon (HIR1 KKRR). Scale bar= 20 µm for confocal and 5 µm for TIRF images **B.** Quantification of the Coefficient of Variation (CV) along a 10µm line of the HIR1 reporter construct transiently expressed in *Nicotiana benthamiana*. Statistical significance is based on pairwise Wilcoxon test (n ≥ 31). **C.** Confocal microscopy images of Nicotiana leaf epidermal cells transiently expressing p35S:HIR1-mNEON or p35S:HIR1(K18Q;K23Q;R50Q)-mNeon (HIR1 KKR) together with compartment marker for Late Endosome/Pre-Vacuolar Compartment (LE/PVC) (W7R), *trans*-Golgi Network/Early Endosome (TGN/EE) (W13R) or Golgi (W18R and W127R). Scale bar= 20 µm **D.** Pearsson’s coefficient was calculated for the number of ROIs n ≥ 30. Statistical significance is based on a pairwise Wilcoxon test.

**Figure S9.**
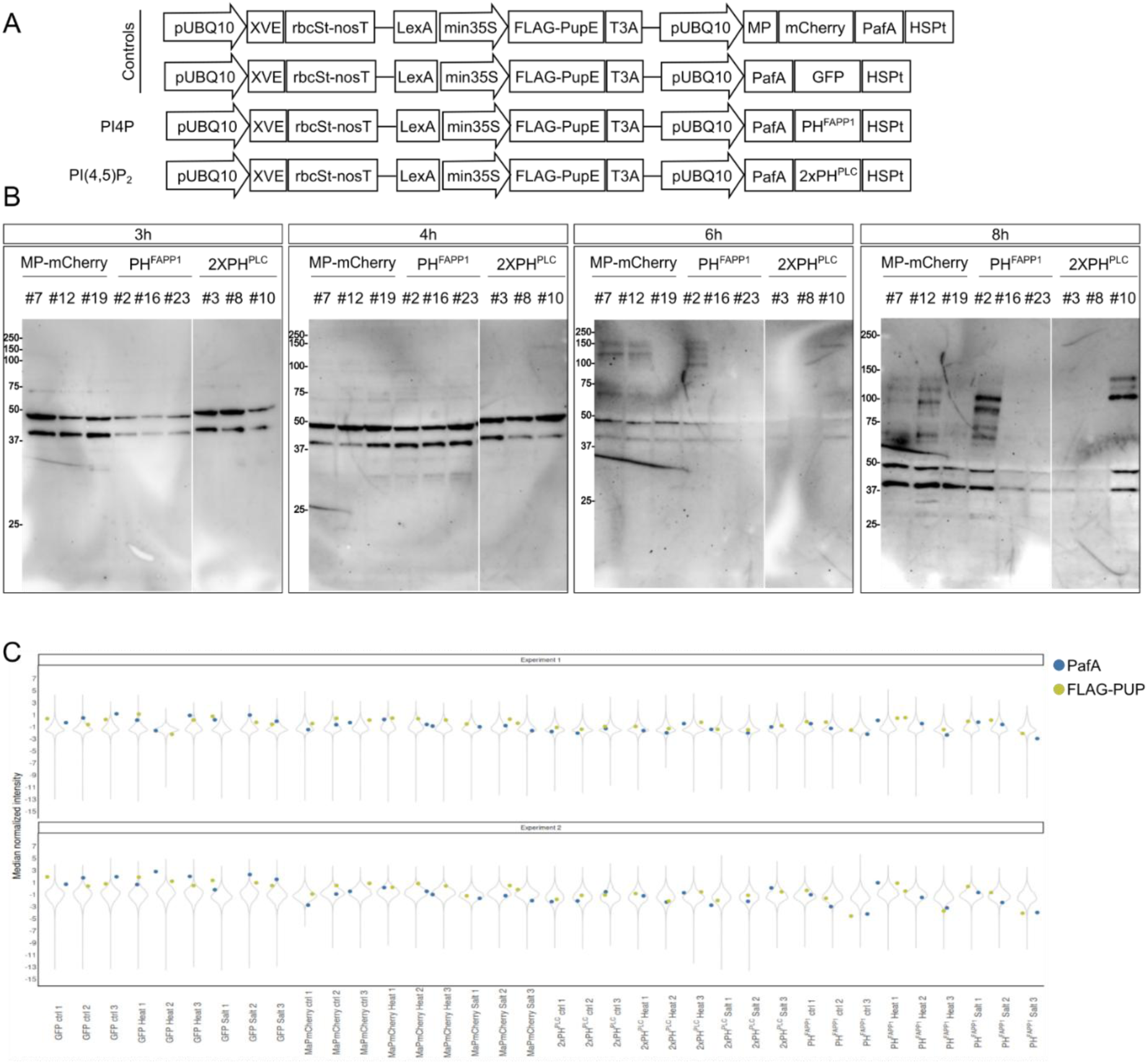
Validation of the inducible PUP-IT constructs used to identify the PIP-associated proteomes in A. thaliana upon stresses. **A.** Schematic representation of the inducible PUP-IT constructs used in *A. thaliana*. For each construct, XVE expression is driven by a UBQ10 promoter and terminated by rbSt and nos terminators. When bound to estradiol, XVE binds the LexA sequence upstream of the min35S promoter and induces the expression of FLAG-PupE. The bait expression is driven by a UBQ10 promoter and terminated by an HSP terminator. **B.** Anti-FLAG western blot of PUP-IT constructs expressed in *A. thaliana* seedlings. Crude lysate from three independent lines at 3, 4, 6, and 8 h after estradiol application. **C.** Median normalized intensity of all proteins for each sample of estradiol-inducible PUP-IT constructs. PafA and FLAG-PUP are highlighted in blue and green, respectively.

**Figure S10.**
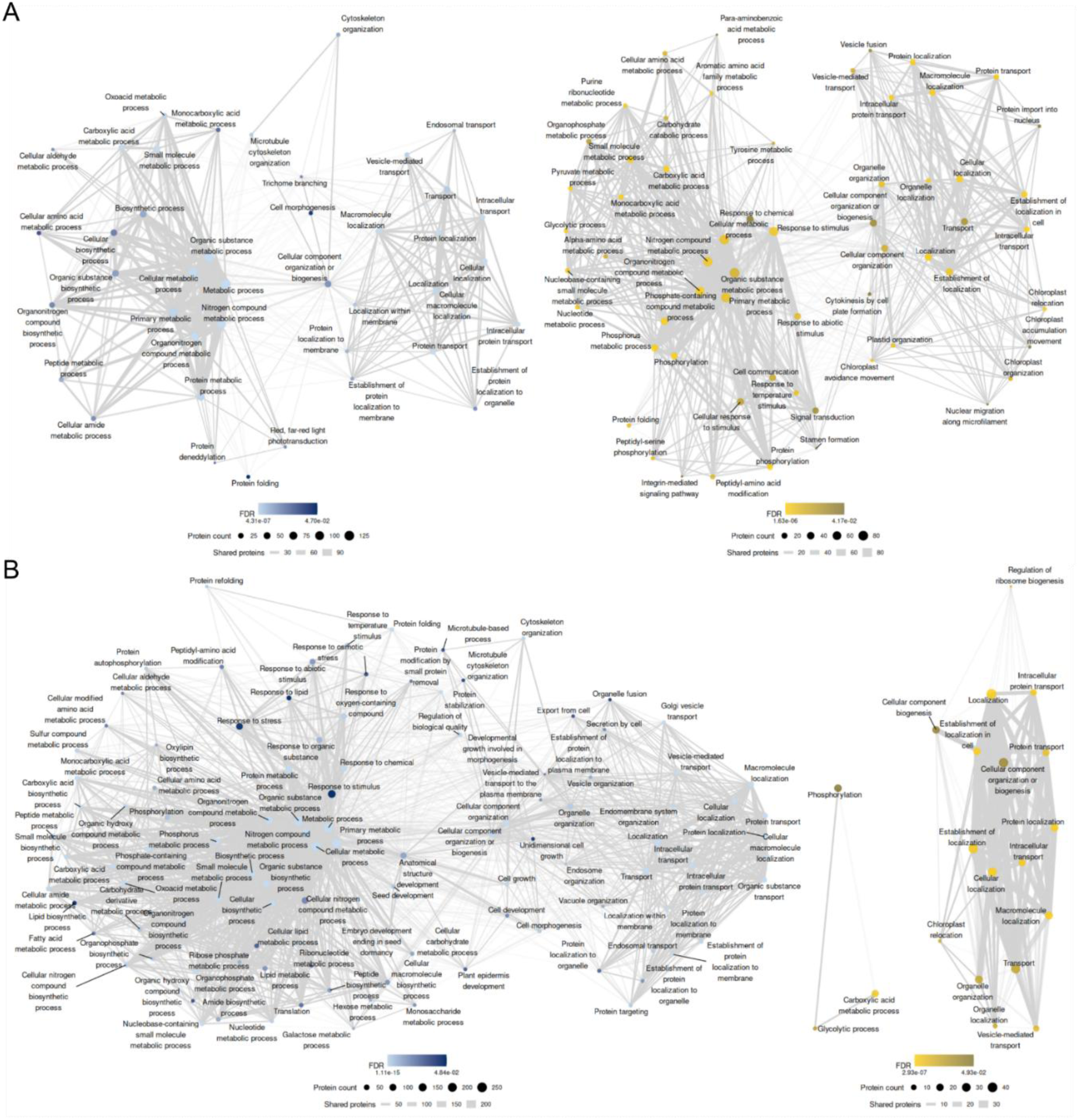
Details of the Gene Ontology analysis of PIP-associated proteomes in A. thaliana upon stresses A,B. Gene Ontology analysis of cellular processes for proteins enriched in salt stress (C) and heat stress (H) compared to control conditions for PH^FAPP1^ in blue and 2xPH^PLC^ in yellow.

**Figure S11.**
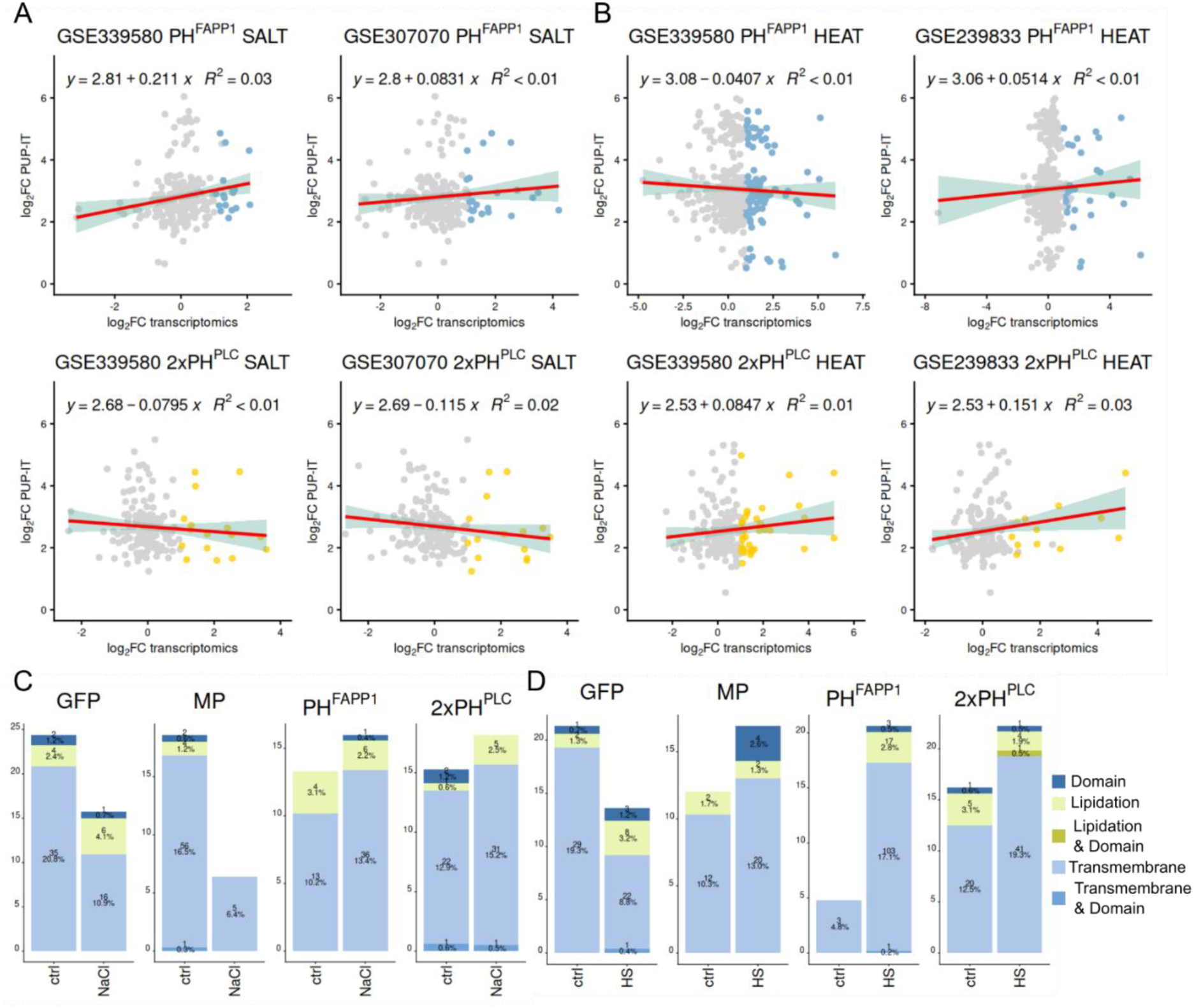
Analyses of proteins enriched in the PIP-associated proteomes in A. thaliana upon stresses A,B. Comparison of PUP-IT data and available transcriptomic data upon salt (A) or heat (B) stress. Linear trend lines with confidence intervals and R-squared values are indicated. Proteins displaying a log_2_FC>1 and an adjusted p-value<0.05 in transcriptomics are shown in color. **C,D.** Proportion of proteins harboring a lipidation site, a lipid binding domain and/or a transmembrane domain for proteins enriched in control, salt stress (NaCl) or heat stress (HS). The number of proteins is indicated in each bar.

---

**Table Supp 1.** N. benthamiana PIP-related proteins identified in the PI4P-associated proteome.

| Function | Protein | Homolog protein in Arabidopsis | Homolog gene in Arabidopsis | Association with PIPs | References |
|---|---|---|---|---|---|
| Vesicular trafficking | NbD019309.1 | SEC11 | AT1G12360 | Subunits of the exocyst complex. In Arabidopsis, the exocyst complex is targeted to the membrane through the interaction of the EXO70 subunit with anionic lipids (including PI4P) | Synek et al., 2021 |
|  | NbD035312.1 | SEC10 | AT5G12370 |  |  |
|  | NbD016283.1 | SNX4 | AT2G15900 | PX-domain-containing protein involved in autophagy and recruited to membranes in a PI3P dependent manner. | Ravussin et al., 2021 |
|  | NbD039878.1 | PICALM4a | AT2G25430 | Phosphatidylinositol-binding clathrin assembly proteins. ANTH domain-containing clathrin adaptor protein | Nguyen et al., 2018 |
|  | NbD038093.1 | ARFC1 | AT3G22950 | ADP-ribosyl factor (ARF)-Like protein that forms a complex with Golgin. In animal cells, recruitment of the complex to Golgi requires interaction with negative membrane lipids. | Lu et al., 2006 |
|  | NbD008379.1 | ARFGAP6 | AT3G53710 | Interact with PI4P at TGN/EE through +ALPS motif | Platre et al., 2018 |
|  | NbE44070901.1 | ARFGAP6 | AT3G53710 |  |  |
|  | NbD047074.1 | ARFGAP8 | AT4G17890 | Predicted ALPS domain. Involved in the recruitment of ARF1 at the Golgi apparatus. | Min et al., 2012 |
|  | NbD028335.1 | SH3P3 | AT4G18060 | SH3P proteins contain a BAR domain. In SH3P2, it mediates its targeting to the PM by interacting with PI(4,5)P<sub>2</sub>. | Doumane et al., 2021 |
| Cytoskeleton/ Membrane contact sites | NbD044616.1 | IQD2 | AT5G03040 | microtubule-binding KLRC-family proteins (CMU1) and IQD2 interact with NET3C which localizes to the PM through a C-terminal lipid-binding motif and interacts with ER-integral VAP27 to form EPCS. | Stefano et al., 2018; Wang et al., 2014 |
|  | NbD049853.1 | CMU1 | AT4G10840 |  |  |
| Membrane organization | NbD042540.1 | REM1.4 | AT5G23750 | REM1.3 is binding phosphoinositides via REM-C anchor | Gronnier et al., 2017 |
|  | NbD051015.1 | REM1.4 | AT5G23750 |  |  |
| Signaling | NbD009807.1 | TMK1 | AT1G66150 | TMK1 is regulated by MAKR proteins which localizes to the plasma membrane in a PI4P dependent manner during gravitropic response. | Marquès-Bueno et al., 2021 |
|  | NbD003518.1 | TMK4 | AT3G23750 |  |  |
|  | NbD001919.1 | PHOT1 | AT3G45780 | RPT2 and NPH3 regulate PHOT activity by phosphorylation in response to light signals. NPH3 relies on electrostatic interactions mediated by PI4P to localizes at the PM and associates with PHOT1 | Reuter et al., 2021 |
|  | NbD009361.1 | PHOT1 | AT3G45780 |  |  |
|  | NbD020628.1 | PHOT2 | AT5G58140 |  |  |
|  | NbD036886.1 | RPT2 | AT2G30520 |  |  |
|  | NbD024504.1 | RIN4 | AT3G25070 | Interactor of the immune receptor RPM1 which specifically interact with PI4P at the plasma membrane through CC domain | Saile et al., 2021 |
|  | NbD039814.1 | RIN4 | AT3G25070 |  |  |
| Protein turnover | NbD031046.1 | OTU11 | AT3G22260 | Deubiquitinating enzyme binding to phospholipids through a polybasic motif. Activity is stimulated by binding to PIPs which can stimulate the endosomal degradation of protein cargo including PINs. | Vogel et al., 2022 |
| Auxin transport | NbD009936.1 | PIN3 | AT1G70940 | PIN protein asymmetric cellular distribution is directly regulated by PINOID or D6PK which both rely on PI4P for targeting PM. | Simon et al., 2016; Barbosa et al., 2016 |

**Table Supp 2.**
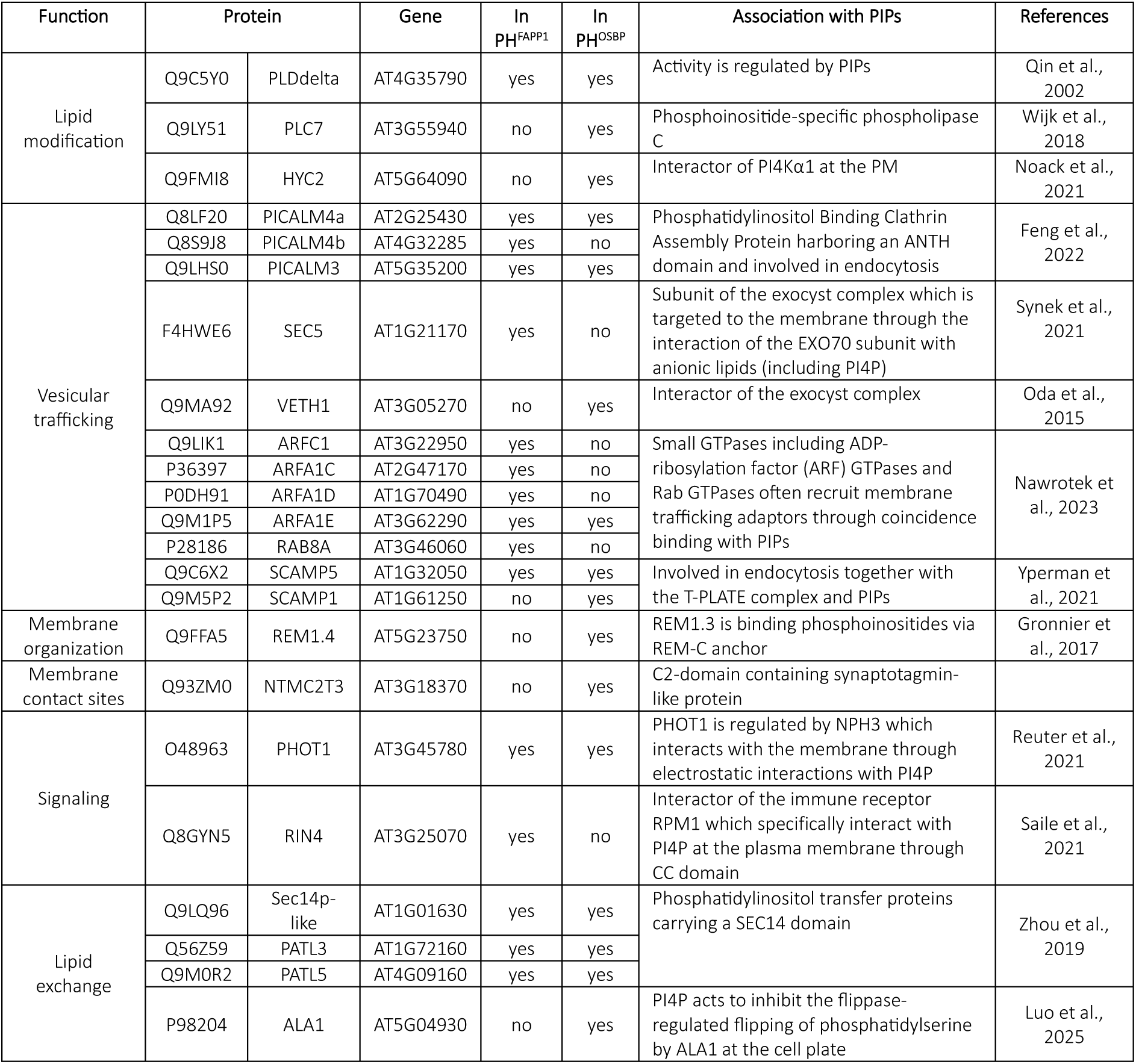
A. thaliana PIP-related proteins identified in the PI4P-associated proteome.

**Table Supp 3.** A. thaliana PIP-related proteins identified in the PI(4,5)P_2_-associated proteome.

| Function | protein |  | gene | Association with PIPs | References |
|---|---|---|---|---|---|
| Lipid modification | Q9C5Y0 | PLDdelta | AT4G35790 | Activity is regulated by PIPs | Qin et al., 2002 |
| Vesicular trafficking | O23140 | AP2µ | AT5G46630 | Recruited to the PM by PI(4,5)P<sub>2</sub> | Doumane et al., 2021 |
|  | Q8VZU2 | SYP132 | AT5G08080 | PIPs can regulate vesicle fusion mediated by syntaxin in eukaryotic organisms | Brillada et al., 2018 |
|  | Q9SF29 | SYP71 | AT3G09740 |  |  |
| Signaling | Q9LK43 | TMK4 | AT3G23750 | TMK1 is regulated by MAKR proteins which localizes to the plasma membrane in a PI4P dependent manner during gravitropic response. | Marquès-Bueno et al., 2021 |
| Membrane organization | Q9M2D8 | Rem1.2 | AT3G61260 | REM1.3 is binding phosphoinositides via REM-C anchor | Gronnier et al., 2017 |

**Table Supp 4.** Arabidopsis transgenic lines used in the study.

| DNA vector | Source |
|---|---|
| 35S:FLAG_PUP(E)_pUBQ10:PafA-eGFP | Zheng et al., 2025 |
| pUBQ10-XVE:FLAG_PUP(E)_mini35S:PafA-eGFP | Zheng et al., 2025 |
| 35S:FLAG_PUP(E)_pUBQ10:MP-mCherry-PafA | This study |
| pUBQ10-XVE:FLAG_PUP(E)_mini35S:MP-mCherry-PafA | This study |
| 35S:FLAG_PUP(E)_pUBQ10:PafA-PH<sup>FAPP1</sup> E50A | This study |
| pUBQ10-XVE:FLAG_PUP(E)_mini35S:PafA-PH<sup>FAPP1</sup> E50A | This study |
| 35S:FLAG_PUP(E)_pUBQ10:PafA-PH<sup>OSBP</sup> | This study |
| 35S:FLAG_PUP(E)_pUBQ10:PafA-1xPH<sup>PLC</sup> | This study |
| pUBQ10-XVE:FLAG_PUP(E)_mini35S:PafA-2xPH<sup>PLC</sup> | This study |
| 35S:FLAG_PUP(E)_pUBQ10:PafA-TubbyC | This study |
| 35S:FLAG_PUP(E)_pUBQ10:PafA-PIP5K2 | This study |
| pUB10:GPA1-mNEON-Ubter-BastaR | This study |
| pUB10:mNEON-PATL5-Ubter-BastaR | This study |
| pUB10:CPK27-mNEON-Ubter-BastaR | This study |
| pUBQ10:BIR2-mScarlet3-Ubter-HygroR | This study |
| pUB10:mNEON-ARL1-Ubter-BastaR | This study |
| pUB10:mNeon-ARL1_Ct-Ubter-BastaR | This study |
| pUB10:mNeon-ARL1_alphaCt-Ubter-BastaR | This study |
| pUB10:ARL1_Nt-mNEON-Ubter-BastaR | This study |
| p35S:HIR1-mNEON-HSPter-BastaR | This study |
| p35S:HIR1(KK)-mNEON-HSPter-BastaR | This study |
| p35S:HIR1(RR)-mNEON-HSPter-BastaR | This study |
| p35S:HIR1(3Q)-mNEON-HSPter-BastaR | This study |
| p35S:HIR1(4Q)-mNEON-HSPter-BastaR | This study |
| UBQ10_XVE:amiRNA_PI4Kα1 | Noack et al., 2021 |
| *pi4kβ1pi4kβ2* | Lin et al., 2019 |
| p35S:eGFP-Lti6b | Kurup et al., 2005 |

**Table Supp 5.** DNA vectors used in the study.

| DNA vector | Source |
|---|---|
| Topo vector | Zheng et al., 2025 |
| pMDC32 | Zheng et al., 2025 |
| pFM | Zheng et al., 2025 |
| MP-mCherry/TOPO | This study |
| PH<sup>FAPP1</sup> E50A/TOPO | This study |
| PH<sup>OSBP</sup>/TOPO | This study |
| 1xPH<sup>PLC</sup>/TOPO | This study |
| 2xPH<sup>PLC</sup>/TOPO | This study |
| TubbyC/TOPO | This study |
| PIP5K2/TOPO | This study |
| PafA-eGFP/pMDC32 | Zheng et al., 2025 |
| PafA-eGFP/pFM | Zheng et al., 2025 |
| MP-mCherry-PafA/pMDC32 | This study |
| MP-mCherry-PafA/pFM | This study |
| PafA-PH<sup>FAPP1</sup> E50A/pMDC32 | This study |
| PafA-PH<sup>FAPP1</sup> E50A/FM | This study |
| PafA-PH<sup>OSBP</sup>/pMDC32 | This study |
| PafA-1xPH<sup>PLC</sup>/pMDC32 | This study |
| PafA-2xPH<sup>PLC</sup>/pFM | This study |
| PafA-TubbyC/pMDC32 | This study |
| PafA-PIP5K2/pMDC32 | This study |
| pGGB000 | Lampropoulos et al., 2013 |
| pGGC000 | Lampropoulos et al., 2013 |
| pGGD000 | Lampropoulos et al., 2013 |
| pGGE000 | Lampropoulos et al., 2013 |
| pGGZ01 | Lampropoulos et al., 2013 |
| HIR1/pGGC | This study |
| HIR1(KK)/pGGC | This study |
| HIR1(RR)/pGGC | This study |
| HIR1(KKR)/pGGC | This study |
| HIR1(KKRR)/pGGC | This study |
| CPK27/pGGC | This study |
| PATL5/pGGC | This study |
| BIR2/pGGC | This study |
| GPA1/pGGC | This study |
| ARG1/pGGC | This study |
| ARL1/pGGC | This study |
| ARL1_Nt/pCCC | This study |
| ARL1_alpha_Ct/pGGC | This study |
| ARL1_Ct/pGGC | This study |
| HSP/pGGE | This study |
| mNEON/pGGD | This study |
| mNEON/pGGB | This study |
| linker-mSc3/pGGD | This study |
| pGGA004 (35S (Cauliflower mosaic virus 35S) promoter) | Lampropoulos et al., 2013 |
| pGGA006 (UBQ10 (UBIQUITIN10) promoter) | Lampropoulos et al., 2013 |
| pGGB003 (B-dummy) | Lampropoulos et al., 2013 |
| pGGD002 (D-dummy) | Lampropoulos et al., 2013 |
| pGGE009 (UBQ10 terminator) | Lampropoulos et al., 2013 |
| pGGF002 (p35S BastaR:t35S) | Lampropoulos et al., 2013 |
| PGGF005 (pUBQ10:Hygro:tOCS) | Lampropoulos et al., 2013 |
| pUB10:GPA1-mNEON-Ubter-BastaR/pZ01 | This study |
| pUB10:mNEON-PATL5-Ubter-BastaR/pZ01 | This study |
| pUB10:CPK27-mNEON-Ubter-BastaR/pZ01 | This study |
| pUBQ10:BIR2-mScarlet3-Ubter-HygroR/pZ01 | This study |
| pUB10:mNEON-ARL1-Ubter-BastaR/pZ01 | This study |
| pUB10:mNeon-ARL1_Ct-Ubter-BastaR/pZ01 | This study |
| pUB10:mNeon-ARL1_alphaCt-Ubter-BastaR/pZ01 | This study |
| pUB10:ARL1_Nt-mNEON-Ubter-BastaR/pZ01 | This study |
| p35S:HIR1-mNEON-HSPter-BastaR/pZ01 | This study |
| p35S:HIR1(KK)-mNEON-HSPter-BastaR/pZ01 | This study |
| p35S:HIR1(R)-mNEON-HSPter-BastaR/pZ01 | This study |
| p35S:HIR1(RR)-mNEON-HSPter-BastaR/pZ01 | This study |
| p35S:HIR1(3Q)-mNEON-HSPter-BastaR/pZ01 | This study |
| p35S:HIR1(4Q)-mNEON-HSPter-BastaR/pZ01 | This study |
| W7R (RabF2a) | Geldner et al., 2009 |
| W13R (VTI12) | Geldner et al., 2009 |
| W18R (Got1p homolog ) | Geldner et al., 2009 |
| W127R (MEMB12) | Geldner et al., 2009 |

**Table Supp 6.**
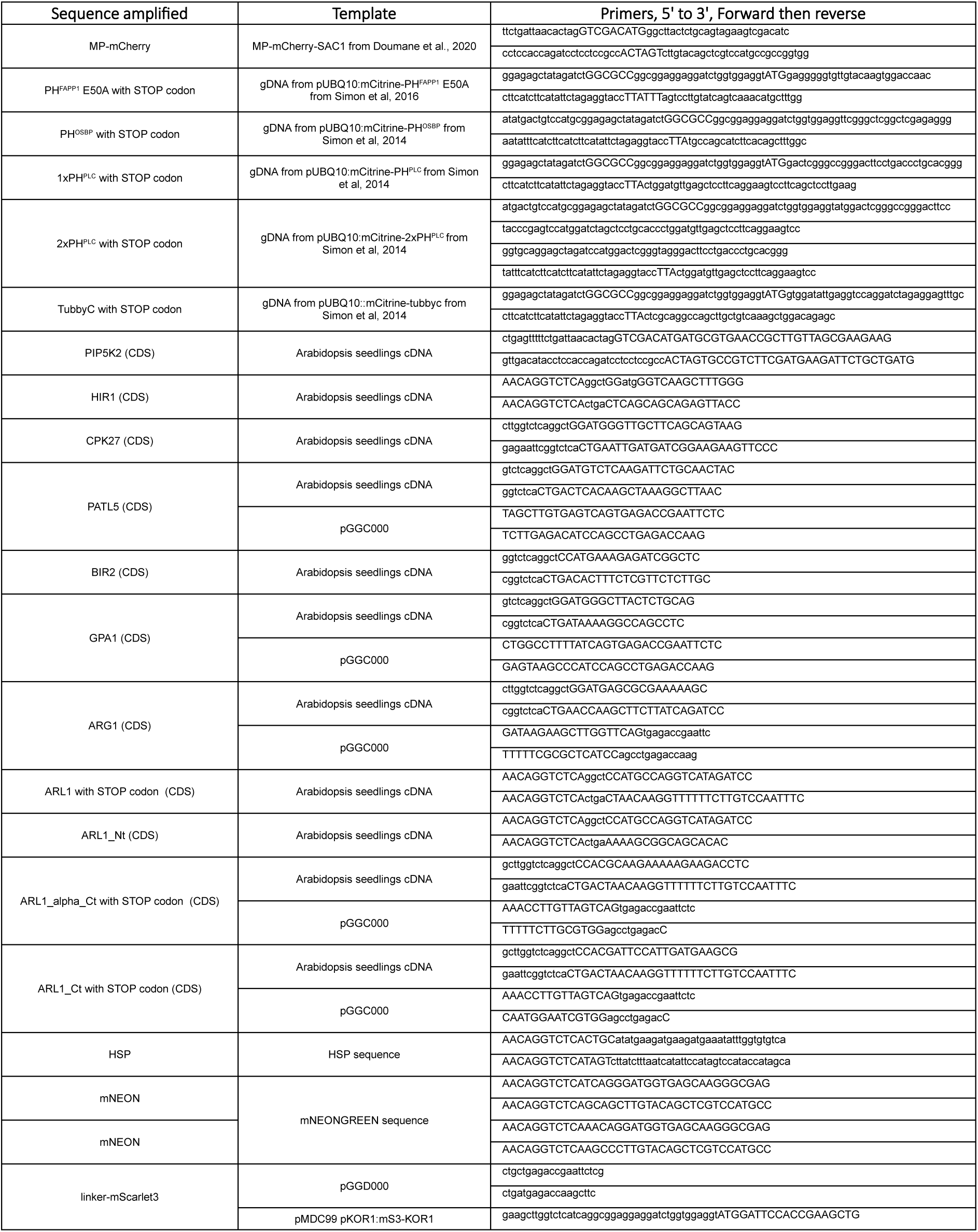
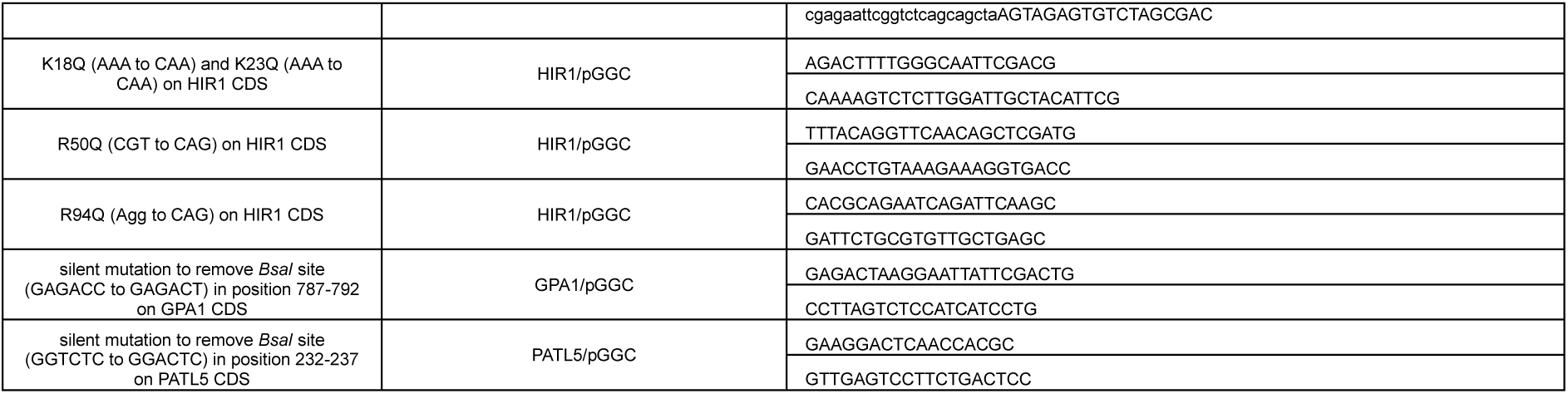
Primers used in the study for cloning.

